# *osr1* maintains renal progenitors and regulates podocyte development by promoting *wnt2ba* through antagonism of *hand2*

**DOI:** 10.1101/2020.12.21.423845

**Authors:** Bridgette E. Drummond, Brooke E. Chambers, Hannah M. Wesselman, Marisa N. Ulrich, Gary F. Gerlach, Paul T. Kroeger, Ignaty Leshchiner, Wolfram Goessling, Rebecca A. Wingert

**Affiliations:** Department of Biological Sciences, Center for Stem Cells and Regenerative Medicine, Center for Zebrafish Research, University of Notre Dame, Notre Dame, 46556, USA; Brigham and Women’s Hospital, Genetics and Gastroenterology Division, Harvard Medical School, Harvard Stem Cell Institute, Boston, MA 02215, USA

**Keywords:** zebrafish, kidney, nephron, podocyte, segmentation, *osr1*, *wnt2ba*, *hand2*

## Abstract

Knowledge about the genetic pathways that control renal cell lineage development is essential to better understand the basis of congenital malformations of the kidney and design regenerative medicine therapies. The embryonic zebrafish kidney, or pronephros, contains two nephrons that are conserved with humans. Recently, the transcription factors Osr1 and Hand2 were found to exert antagonistic influences to balance kidney specification (Perens et al., 2016). Here, we performed a forward genetic screen in zebrafish to identify nephrogenesis regulators, where whole genome sequencing of the novel *oceanside* (*ocn*) mutant revealed a nonsense mutation in *osr1. ocn* mutants evince severe pronephros defects including abrogation of podocytes and proximal tubule cells. Our studies reveal that *osr1* is not needed to specify renal progenitors, but rather required to maintain their survival. Additionally, *osr1* is requisite for expression of the canonical Wnt ligand *wnt2ba*, where *wnt2ba* is expressed in the intermediate mesoderm (IM) and later restricts to podocytes. Deficiency of *wnt2ba* reduced podocyte progenitors, where overexpression of *wnt2ba* was sufficient to rescue the podocyte lineage as well as *osr1* loss of function. Finally, we demonstrate that reciprocal antagonism between *osr1* and *hand2* mediates podocyte development specifically by controlling *wnt2ba* expression in the IM. Together, our data show that Osr1 is essential for a sequence of temporal functions that mediate the survival and lineage decisions of IM progenitors, and subsequently the maintenance of podocytes and proximal tubule epithelium in the embryonic nephron.

## INTRODUCTION

The kidney is the organ that cleanses our blood and initiates the process of waste excretion. The portion of the kidney that makes this possible are the nephrons, which are composed of a blood filter, tubule and collecting duct. The blood filter itself is composed of several cellular features including a capillary bed with a fenestrated endothelium, known as the glomerulus, and the space that the glomerulus is housed in, known as the Bowman’s capsule. Octopus-like epithelial cells known as podocytes are situated in opposition to a specialized glomerular basement membrane (GBM) surrounding the capillaries (Ichimura et al. 2017). Filtration is accomplished due to the layered ultrastructure of fenestrated epithelium, GBM, and podocytes, which keeps large bulky particles from entering the tubule (Grahammer 2017; Pavenstadt et al. 2003). Podocytes form elaborate cellular extensions and are connected to adjacent podocytes through cell membrane based protein interactions that create a specialized barrier known as the slit diaphragm (Grahammer 2017). The slit diaphragm does allow small or appropriately charged molecules to pass, which initiates a filtrate product that flows into the tubule and is subsequently augmented by specialized solute transporters arranged in a segmental pattern to create a concentrated waste product (Ichimura et al. 2017; Garg 2018). Within the nephron tubule, the proximal segments perform the bulk of reabsorption, particularly of organic molecules, while the distal segments fine tune the amount of water within the filtrate (Zhuo and Li, 2013). The two kidneys in our body, each composed of around a million nephrons, filter all of the blood in our body almost 30 times daily to produce 1-2 quarts of urine (NIDDK). Any damage or deficit of the specialized cells of the kidney is detrimental to this process, as the human kidney has a limited regenerative capacity. Furthermore, there are currently no therapeutic interventions that can reverse the damage to the kidneys for patients with acquired kidney diseases and birth defects (Wiggins 2007; Romagnani et al. 2017; Reiser and Sever 2013). One reason for this is our limited understanding about kidney developmental pathways.

Early in embryogenesis, the intermediate mesoderm (IM) gives rise to the earliest form of the kidney, known as the pronephros. The paraxial mesoderm (PM) and lateral plate mesoderm (LPM) fields flank the developing IM (Gerlach and Wingert 2013; Perens et al. 2016). While functional in lower vertebrates, the pronephros is vestigial in mammals (Little and McMahon 2012). This structure degenerates to give rise to the mesonephros, which is the terminal kidney in fish and amphibians, but in mammals it is followed by the metanephros (McMahon 2016). Of note, while many organs such as the brain continue to develop post-gestation, humans are born with metanephric kidneys with a static number of nephrons (Luyckx et al. 2011; Little 2016). However, this is not the case for teleosts such as the zebrafish (*Danio rerio*) that continue to grow nephrons throughout their lifetime and are even capable of neonephrogenesis upon injury (Drummond and Wingert 2016; Drummond and Davidson 2016).

Despite these differences, zebrafish do exhibit fundamental genetic and morphological similarities in kidney organogenesis to mammals. For example, the mammalian renal progenitor markers *Lim homeobox 1* (*LHX1)* and *paired box gene 2 (PAX2)* are orthologous to *LIM homeobox 1a* (*lhx1a*) and *paired box 2a* (*pax2a*), which are also renal progenitor markers in zebrafish (Diep et al. 2011, Naylor et al. 2013). Additionally, zebrafish podocytes morphologically resemble mammalian podocytes, and express a suite of markers including *Wilms tumor 1a, Wilms tumor 1b, nephrosis 1, congenital Finnish type (nephrin), nephrosis 2, idiopathic, steroid-resistant (podocin)* (*wt1a/b, nphs1*, and *nphs2)* that closely correspond to the human homologs *WT1, NPHS1, and NPHS2*, respectfully (Hsu et al. 2003; Bollig et al. 2006; O’Brien et al., 2011; Zhu et al. 2016). Further, the zebrafish nephron exhibits a conserved collection of solute transporter genes that have been shown to be arranged into two proximal and two distal segments similar to other vertebrates including mammals (Figure 1A) (Wingert et al. 2007; Wingert et al. 2011; Desgrange and Cereghini 2015). The genetic conservation combined with the simplicity of the two-nephron and single blood filter pronephros makes the embryonic zebrafish kidney an accessible and powerful genetic model to gain insight into the many puzzles and complexities of kidney development.

**Figure 1:**
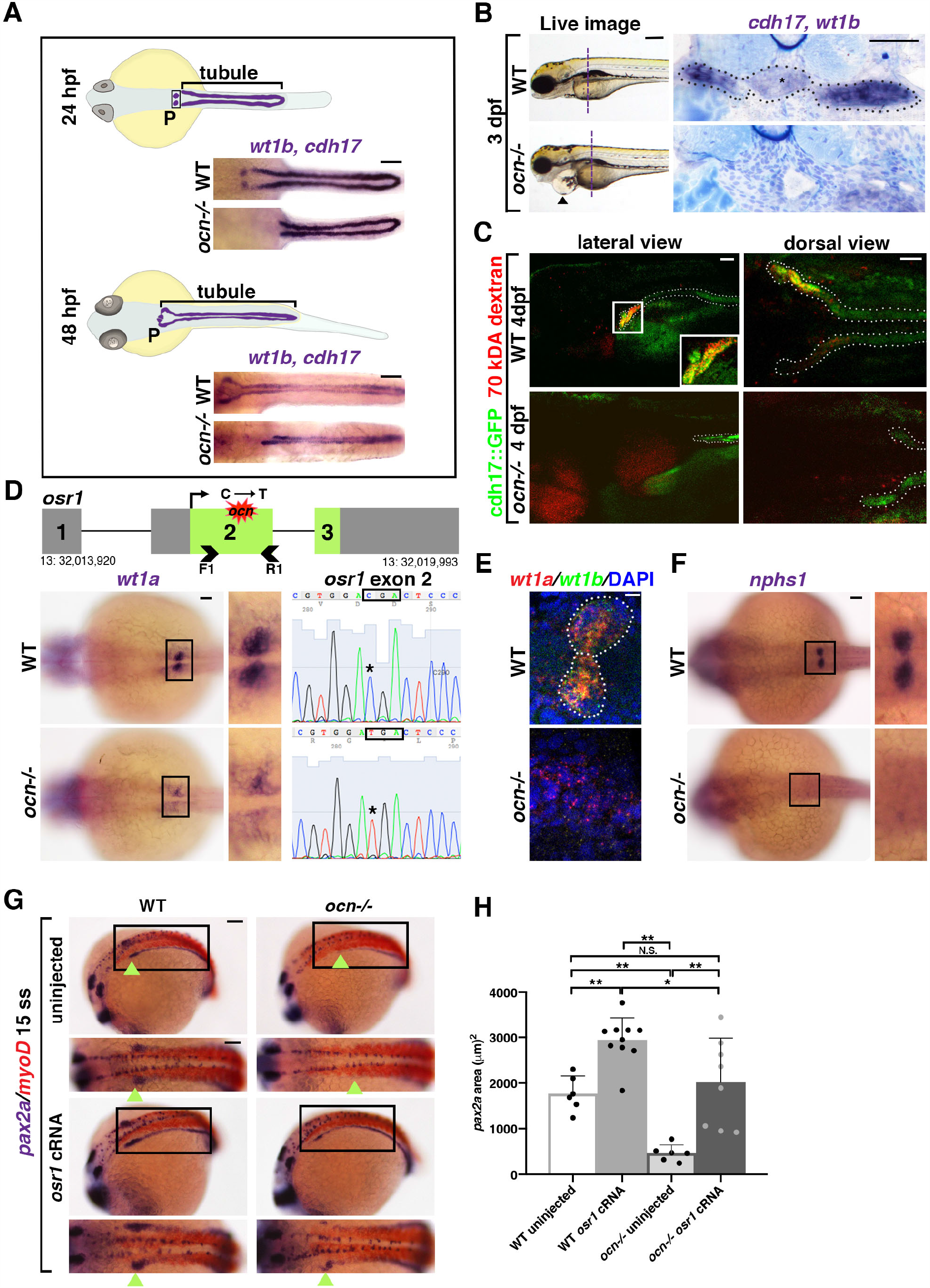
The ENU mutant *ocn* has a proximally abrogated pronephros due to a lesion in *osr1*. **A**. At 24 hpf, the zebrafish pronephros contains two clusters of podocytes and two nephron tubules. By 48 hpf, the pronephros is functional as the podocyte progenitors have migrated to the midline and fused. In *ocn-/-*mutants, podocyte progenitors (*wt1b*) are reduced at both stages. The pronephric tubules (*cdh17*) were truncated at 24 hpf, which became more dramatic at 48 hpf. Scale bar is 50 µm. **B**. A live time course of *ocn* revealed pericardial edema beginning at 72 hpf, as indicated by black arrow heads. This fluid imbalance was symptomatic of organ dysfunction. Scale bar is 70 µm. WISH experiments to view podocytes and tubule (*wt1b, cdh17*) were also conducted at 72 hpf. JB-4 serial sectioning was conducted on three WT and three *ocn-/-*embryos to examine the anterior pronephros, the location is marked by the dashed vertical line. WT siblings had an intact pronephros (dotted outline), including a glomerulus (asterisk) with two tubules. Mutant sections of this same region had no discernable blood filter or tubule structure. Scale bar is 50 µm. **C**. At 48 hpf, *ocn*::cdh17::GFP embryos were injected with 70 kDA rhodamine dextran (red). These embryos were assessed at 96 hpf. Nephron tubules are shown by the dotted outline. WT siblings exhibited no edema and appeared to uptake the dextran in the proximal region, as indicated by yellow coloration (inset). However, in mutants with pericardial edema and truncated tubules, there was no evidence of dextran within the tubule, suggesting that active uptake was not occurring in these mutants. Scale bar is 15 µm for lateral images and 50 µm for dorsal views. **D**. After assessment of the genetic candidates obtained via whole genome sequencing, *osr1* appeared to be an attractive possibility due to a C to T SNP that was predicted to cause a premature stop codon. The predicted lesion (red shape) is located in exon 2 of *osr1*. We designed primers that flanked exon 2 (arrow heads) for Sanger sequencing. Embryos with reduced *wt1a* WISH staining exhibited a “TGA” codon within exon 2 of *osr1* that is normally a “CGA” codon in WT embryos. Scale bar is 30 µm. **E**. To confirm that *wt1a*+ podocytes were reduced in *ocn-/-*, FISH with *wt1a* and *wt1b* was performed at 24 hpf. There were little to no double positive cells seen in genotype-confirmed mutants, whereas both clusters of *wt1a/b*+ podocytes were evident in WTs. Scale bar is 10 µm. **F**. The slit diaphragm marker *nphs1* was similarly reduced in 24 hpf *ocn* mutants. Scale bar is 50 µm. **G**,**H**. At the 15 ss, *pax2a* marks the developing IM, the beginning of which is shown with green arrowheads. In *ocn-/-*, the anterior region of *pax2a* is decreased. When *ocn-/-*was injected with *osr1* cRNA, *pax2a* expression was restored. Interestingly, *pax2a* was significantly expanded in WT embryos injected with *osr1*. Absolute area measurements of *pax2a* were taken from somites 1-5. P-values: **p<0.001, *p<0.05, N.S. = not significant. Scale bar is 50 µm.

A critical regulator of kidney development in both zebrafish and mammals is the zinc-finger transcription factor *odd skipped-related 1* (*OSR1*). In mice, *Osr1* is one of the earliest markers of the IM and fate-mapping studies have shown that *Osr1*+ cells differentiate into renal progenitors and renal-associated vasculature (Mugford et al. 2008). *Osr1-/-*mice fail to express renal progenitors or develop metanephric kidneys, which contributes to embryonic lethality (James et al. 2006; Wang et al. 2005). Similar to mouse studies, zebrafish *osr1* is an initial marker of the IM (Mugford et al. 2008). Further, knockdown of *osr1* causes edema, disrupts glomerular morphogenesis, and reduces proximal tubule in both zebrafish and *Xenopus* (Tena et al. 2007). Subsequent studies have confirmed these findings (Mudumana et al. 2008; Tomar et al. 2014; Neto et al. 2012; Perens et al. 2016; Perens et al. 2020, bioRxiv), though the intriguing observation that *osr1* knockdown causes the kidney structure to be lost in the region abutting somites 3-5 is not fully understood. In humans, mutations in *OSR1* have been clinically linked to hypomorphic kidneys, making the continued study of this factor and its genetic regulatory network a necessity (Zhang et al. 2011).

In this study, we report the discovery of the zebrafish *oceanside* (*ocn*) mutation, which was identified from a forward genetic screen for defects in kidney development based on a striking reduction in podocytes and anterior pronephros tubule (Kroeger et al. 2014; Kroeger et al. 2017; Chambers et al. 2018). Whole genome sequencing revealed the causative lesion of this mutant was a premature stop codon in exon 2 of *osr1*. In addition to *ocn*-/-recapitulating previously observed alterations in mesoderm-derived tissues, reductions in nephron tubule and podocytes were rescued with ectopic *osr1* cRNA. Interestingly, we found that while *osr1* was not needed to initially establish the renal progenitor field, *osr1* was needed to maintain the renal progenitors, as they underwent apoptosis in the absence of Osr1 activity. We also found that *wnt2ba* transcripts were expressed in podocytes and that this expression was significantly decreased in *ocn-/-*. Loss of *wnt2ba* led to a reduction in podocytes, and ectopic *wnt2ba* was sufficient to partially restore podocyte development in *ocn-/-*. Further, we placed *wnt2ba* downstream of the antagonistic influences exerted by *osr1* and *hand2* during IM ontogeny. Together, these data illuminate novel functions of Osr1, which are essential to forwarding our understanding of renal lineage development and may have important implications for congenital kidney defects and diseases as well.

## RESULTS

### *ocn* encodes a premature stop codon in *osr1* and mutants exhibits defective podocyte and pronephric tubule development

A forward genetic haploid screen was performed to identify regulators of nephrogenesis using the zebrafish pronephros model (Kroeger et al. 2014; Kroeger et al. 2017; Chambers et al. 2018). The *ocn* mutant was isolated due to its loss of podocytes and anterior pronephros (Figure 1A). Whole mount *in situ* hybridization (WISH) was performed to delineate the two pronephros tubules based on expression of transcripts encoding *cadherin 17* (*cdh17*) and the podocytes based on *wt1b* expression at the 24 and 48 hours post fertilization (hpf) stages (Figure 1A). Both tubule length and podocyte area were found to be significantly reduced in *ocn* mutants at these time points compared to wild-type (WT) embryos (Figure 1 - figure supplement 1A). By 72 hpf, *ocn* mutants exhibited dramatic pericardial edema that progressed in severity through 120 hpf, and was ultimately lethal (Figure 1B, Figure 1 - figure supplement 1B). Since the kidneys play a major role in fluid homeostasis, this phenotype was a probable indicator of renal dysfunction.

To explore this further, WISH staining to assess tubule and podocyte morphology was conducted on 72 hpf *ocn* and WT embryos. The animals were embedded in JB-4 plastic resin and serially sectioned. In WT embryos, the blood filter could be detected as a mass of dense capillaries containing glomerular podocytes (*wt1b+*) flanked by *cdh17+* tubules (Figure 1B). In *ocn* mutants, however, both the glomerulus and the proximal tubules were abrogated (Figure 1B). Instead, a dilated dorsal aorta was identified in this region (Figure 1B).

While it was clear that the proximal pronephros was absent in *ocn* mutants, it was uncertain if this truncated kidney retained any functionality. Kidney functionality was assessed using an endocytosis assay whereby 70 kDA rhodamine-dextran was injected into the vasculature of *ocn*::*cdh17*::GFP embryos, which exhibited a pronephric truncation that phenocopied WISH experiments at 3 days post fertilization (dpf) onwards (Figure 1 - figure supplement 1C). Transgenic animals were injected with rhodamine-dextran at 48 hpf and then assessed at 48 hours post injection (hpi). While dextran was endocytosed in the proximal tubule of non-edemic WT siblings, there was no dextran observed in the truncated tubules of the edemic *ocn* mutant embryos (Figure 1C). Additionally, we assessed epithelial polarity through immunofluorescence (IF) staining of Na-K-ATPase, which marks transporters localized along the basolateral sides of kidney epithelial cells, and aPKC, which marks the apical side of these epithelia (Gerlach and Wingert 2014). This experiment revealed a similar reduction in tubule and podocytes in *ocn* as seen with our WISH experiments using the markers *cdh17/wt1b* (Figure 1 - figure supplement 1D). Together, this provided strong evidence that the stunted pronephros in *ocn-/-*was functionally defective.

Next, to identify the causative lesion in *ocn*, whole genome sequencing was conducted on pools of genomic DNA collected from 24 hpf WISH-identified putative mutants and WT siblings (Leschiner et al. 2012; Ryan et al. 2013). Analysis of the sequencing was performed using SNPtrack software, whereby we detected a strong candidate SNP that was centrally located on chromosome 13 (Figure 1 - figure supplement 2A). Specifically, the putative SNP encoded a missense C to T mutation and was predicted to result in an amino acid substitution from an arginine to a premature stop codon in exon 2 of *osr1* (Figure 1D, Figure 1 - figure supplement 2A).

To further assess if the predicted stop codon in exon 2 of *osr1* was linked with the *ocn* phenotype, we performed additional genotyping analysis. For this, genomic DNA was isolated from individual embryos that had been identified as *ocn* mutants or WTs, based on WISH with the podocyte marker *wt1a* at 24 hpf, and PCR was performed to amplify exon 2 of *osr1* followed by direct Sanger sequencing (Figure 1D). Out of 20 *ocn* embryos with reduced *wt1a* staining, all 20 were homozygous for the C to T mutation in exon 2 of *osr1* (Figure 1D). Protein alignment showed that zebrafish and human OSR1 protein are 264 amino acids (aa) and 266 aa in length, respectively (Figure 1 - figure supplement 2B). While they exhibit 77% conservation in overall aa sequence, the three zinc finger domains responsible for DNA binding activity are 100% conserved across humans, mice and zebrafish (Figure 1 - figure supplement 2B). The *osr1* genetic lesion in *ocn-/-*would place a premature stop codon at residue 165 before all three zinc finger domains (Figure 1 - figure supplement 2B). This suggested that the truncated Osr1 protein produced in *ocn-/-*would not contain any functional domains and would thus be unable to act as a targeted transcription factor. Next, we verified the effectiveness of a splice-blocking *osr1* morpholino with RT-PCR (Figure 1 – figure supplement 3). *osr1* morphants had a decrease in podocytes and proximal tubule that phenocopied *ocn-/-*and was consistent with phenotypes reported in previous studies that utilized the same morpholino (Mudumana et al. 2008, Tomar et al. 2014, Neto et al. 2012, Perens et al. 2016) (Figure 1 – figure supplement 3).

Previous literature has indicated that *osr1* acts to restrict venous development in order to promote other mesodermal fates such as the kidney and the pectoral fins (Mudumana et al. 2008; Perens et al. 2016; Neto et al. 2012). At 4 dpf, Alcian blue staining indicated that *ocn-/-*possessed shorter, malformed pectoral fins (Figure 1 - figure supplement 1E). The fin bud area, which gives rise to pectoral fins, was significantly reduced in *ocn-/-*mutants compared to siblings as seen by the marker *MDS1 and EVI1 complex locus* (*mecom*) at 24 hpf (Figure 1 - figure supplement 1F). Additionally, Alcian blue staining revealed altered craniofacial cartilage formation in mutants which fits with previous literature placing *osr1* as a regulator of palatogenesis in zebrafish and mice (Swartz et al. 2011; Liu et al. 2013) (Figure 1 - figure supplement 1E). In sum, these mesodermal phenotypes were consistent with *osr1* deficiency.

Next, we evaluated other aspects of pronephros development. As *wt1a* expression appeared to be severely diminished and also disorganized in *ocn-/-*, we evaluated additional markers to better understand the features of podocyte lineage development in mutants. Podocytes were examined at 24 hpf using a *wt1a/wt1b* double fluorescent *in situ* (FISH). While clusters of *wt1a+wt1b+* podocytes were visible in siblings, mutants had a scarcity of double positive cells (Figure 1E). There was also a dearth in *nphs1+* cells, which is a marker of the podocyte slit diaphragm, and suggested that podocyte differentiation was also disrupted (Figure 1F). Additionally, *ocn-/-*embryos displayed diminished *pax2a* expression at the 15 ss compared to siblings (Figure 1G), again characteristic of *osr1* morphants in previous studies (Mudumana et al. 2008, Neto et al. 2012, Perens et al. 2016).

To test whether the mutation in *osr1* was the specific cause of this phenotype, we performed rescue studies. Injection with *osr1* capped RNA (cRNA) rescued this domain in *ocn* mutants and expanded it in WT siblings (Figure 1G,H). In sum, *ocn* mutants reciprocated a multitude of mesodermal phenotypes seen in *osr1* literature in zebrafish and across taxa. The ability of *osr1* cRNA to rescue key mesodermal phenotypes in *ocn-/-* and the catastrophic nature of the *osr1* mutation, we concluded that Osr1 deficiency is responsible for the *ocn* phenotype.

### Kidney progenitors are specified in *osr1* deficient animals, but subsequently undergo apoptosis

Previous studies suggest that the anterior pronephros abrogation in *osr1* zebrafish morphants is due to a fate change where blood/vasculature and endoderm form instead of renal progenitors (Mudumana et al. 2008; Terashima et al. 2014; Perens et al. 2016). Interestingly, in the *Osr1* mouse knockout model, there was an increase in apoptosis that occurred within the kidney tissue (James et al. 2006). However, in both models, renal progenitors are initially established (James et al. 2006; Mudumana et al. 2008). Thus, we next sought to delineate the cellular dynamics of renal progenitor specification in our *ocn* mutant model, and to address if alterations in proliferation or apoptosis occur during pronephros development in the absence of *osr1*.

To investigate this, we first performed WISH studies. The LPM is marked by *T-cell acute lymphocytic leukemia 1* (*tal1*), and gives rise to hemangioblasts (Gering et al. 1998; Liao et al. 1998). The IM and hemangioblast domains at the 7 ss were not noticeably different between WT and *ocn-/-*embryos, as indicated by the markers *pax2a*, and *tal1* (Figure 2A). However, as previously noted, by the 15 ss there was a decrease in the anterior-most domain of *pax2a* expression in *ocn-/-*embryos (Figure 1G). To further assess the anterior *pax2a*+ cells between the 7 ss and 15 ss, we performed double FISH studies in WT and *ocn-/-*embryos to assess *pax2a* and *tal1* expression. DAPI staining was also utilized to discern features such as the trunk somites, which allowed for accurate staging. Embryos were flat-mounted and imaged as previously described (Figure 2 - figure 2 supplement 2A) (Cheng et al. 2014). Further, IF was also performed on these samples with anti-caspase-3 antibody to assess the number apoptotic bodies or anti-pH3 to identify proliferating cells. In our analysis we focused on the changes to these markers within somites 1-5, as the IM adjacent to somite 3 gives rise to podocytes (O’Brien et al. 2011).

**Figure 2:**
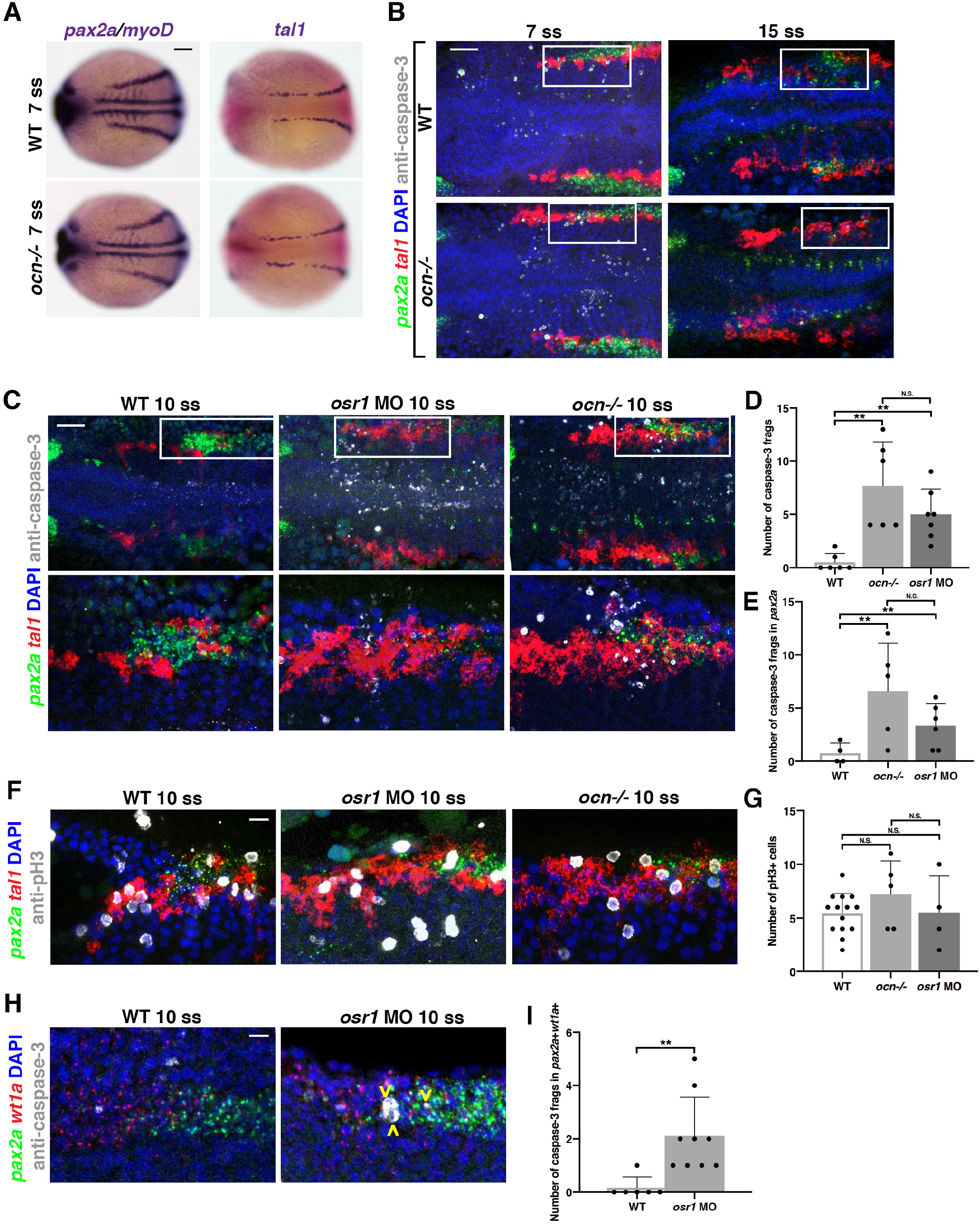
*osr1* is required to maintain and promote kidney development at the expense of hemagioblasts. **A**. Although *pax2a* is restricted in 15 ss mutants, at 7 ss, *ocn-/-*embryos had a *pax2a* domain that appeared to occupy the same domain as WT siblings. Similarly, the hemangioblast marker *tal1* appeared mostly WT in *ocn-/-*at 7 ss. Scale bar is 50 µm **B**. FISH with probes for *pax2a* (green) and *tal1* (red) and ICC using anti-caspase-3 (white) to mark apoptotic cells was conducted at 7 ss and 15 ss. The number of fragments in the combined *pax2a* and *tal1* fields from somites 1-5 were increased in mutants at 7 ss, but not at 15 ss. Scale bar is 35 µm **C**,**D**,**E**. At 10 ss, little to no caspase-3 fragments were seen in *pax2a* or *tal1* domains from somites 1-5, but a significant number were seen in *osr1* morphants and *ocn-/-*. The *tal1* domain was also expanded in both loss of function models. Scale bar is 35 µm. **F**,**G**. ICC with the proliferative cell marker anti-pH3 was also conducted. Despite the expansion in *tal1* in *osr1* deficient models, there was no significant change in the number of proliferating cells. Scale bar is 10 µm. **H**,**I**. FISH experiments were conducted to assess changes in apoptosis in *wt1a+pax2a+* podocyte progenitors. There was a significant increase in the number of apoptotic fragments within this field in mutants compared to WT siblings. Scale bar is 10 µm. A minimum of three individuals were assessed for each group across experiments. Photos are max intensity projections from z-stacks, and each side of mesoderm was quantified individually. P-values: **p<0.001, N.S. = not significant.

Beginning at the 7 ss, *ocn*-/-embryos exhibited a significant increase in the number of caspase-3+ fragments within the combined *tal1* and *pax2a* fields near somites 1-5 (Figure 2B). However, by the 15 ss, few apoptotic fragments were visible in the area of interest, with no significant differences between WT and *ocn-/-*embryos (Figure 2B). Similar to the 7 ss, we found a significant increase in the number of total caspase-3+ fragments at the 10 ss in *ocn-/-*mutants and *osr1* morphants, while the WT siblings had little to no apoptosis occurring in this area (Figure 2C-E). Interestingly, most of the apoptosis that occurred in mutants and morphants happened within the *pax2a* kidney field specifically (Figure 2E). Another finding of note was that the number of caspase-3+ fragments was not significantly different between *osr1* morphants and *ocn-/-*for either assessment (Figure 2D,E).

To further understand the cell dynamics across this time course, absolute area measurements of *pax2a* and *tal1* were taken at 7, 10, and 15 ss from somites 1-5 in WT and *ocn-/-*. Surprisingly, the area of the *tal1* domain was already expanded at 7 ss, and continued to expand through the 15 ss (Figure 2 – figure supplement 1). However, across the three time points examined, a reduction in *pax2a* area was only significantly different between WT and *ocn-/-*at the 15 ss (Figure 2 – figure supplement 1). Further, although the *tal1* field was expanded in *ocn-/-*embryos at the 10 ss, there was no significant difference in proliferating pH3+ cells between WT and *osr1* loss of function models (Figure 2F,G). Additionally, no significant changes in proliferation were seen between mutants and WTs at the 8 ss (Figure 2 – figure supplement 1).

To determine if apoptosis was occurring within podocyte progenitors in the *pax2a* kidney field, we performed an additional FISH with *wt1a* and *pax2a* at 10 ss. During this time point, while *pax2a* expression begins adjacent to somite 3, *wt1a* is expressed from somites 1-3 (Figure 2H). Similar to the *pax2a* domain, the *wt1a* domain did not appear to be reduced at this time point, though it does become restricted and disorganized by 24 hpf (Figure 1D). We found a significant increase in caspase-3+ fragments that were double positive for *wt1a* and *pax2a* in *osr1* morphants compared to WTs (Figure 2H,I). These results demonstrated that abnormal apoptosis occurred in podocyte progenitors due to loss of *osr1*. In sum, *osr1* is not needed to initiate the *pax2a* progenitor pool, but it is needed to maintain this population, including the podocyte precursors, during pronephros development.

### Ectopic *osr1* is transiently sufficient to rescue renal progenitors

Our observation that *pax2a+* renal progenitors arise in *ocn* mutants, but are not maintained, is consistent with previous data that *osr1* knockdown leads to a reduced *pax2a+* renal progenitor field by the 14 ss (Mudumana et al. 2008). As *pax2a* expression marks both podocyte and tubule precursors (O’Brien et al. 2011), we hypothesized that *osr1* is likely needed for podocyte and tubule progenitor maintenance. To determine this, we performed a series of rescue studies in our *ocn* mutants to test if one or both of these compartments requires *osr1* for its maintenance.

First, we tested whether overexpression of *osr1* mRNA was sufficient to rescue podocytes in *ocn* mutants by assessing *wt1b* expression, which specifically marks the podocyte lineage (Bollig et al. 2006; O’Brien et al. 2011). Provision of *osr1* mRNA robustly rescued the development of *wt1b+* podocytes in *ocn* mutants at the 15 ss (Figure 3A,B). However, by the 22 ss, we were only able to achieve a partial podocyte rescue, though tubules within the same individuals appeared to be WT in length (Figure 3C,D). Consistent with this, we were unable to obtain a podocyte rescue at 24 hpf (data not shown), though again we could achieve a rescue of the truncated tubules (Figure 3E,F). Interestingly, overexpression of *osr1* was sufficient to induce ectopic *cdh17+* cells in about 55% of injected embryos (Figure 3G). It should be noted that *osr1* cRNA did lead to a decrease in body axis length when compared to uninjected WTs and mutants, which in turn affected pronephros length measurements (Figure 3 - figure supplement 1A). Despite this, the percentage of kidney length to body length was not significantly different between embryos injected with *osr1* cRNA and uninjected animals.

**Figure 3:**
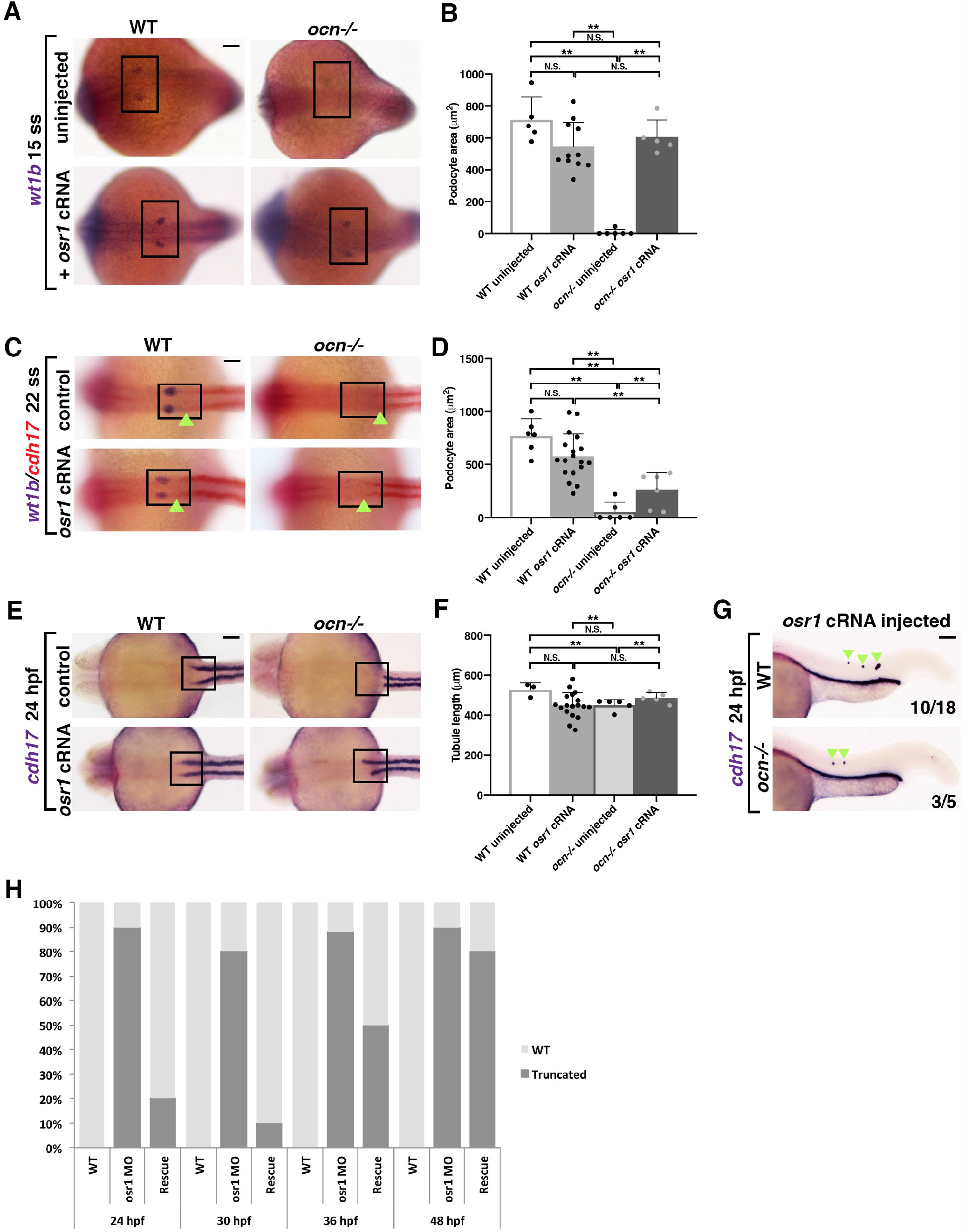
*osr1* is required for continued development of kidney lineages. **A**,**B**. Embryos from *ocn* incrosses were injected at the 1-cell stage with 50 pgs of *osr1* cRNA and examined. At 15 ss, podocytes (*wt1b*) were robustly rescued in *ocn-/-*. **C**,**D**. However, by 22 ss, podocytes were only partially rescued in *ocn-/-*injected embryos, while tubule length in the same animals were not significantly different from WTs, as shown by the green arrowheads. **E**,**F**,**G**. Truncated tubule was still able to be rescued in mutants at 24 hpf. Further, overexpression of *osr1* induced ectopic tubule formation (green arrowheads). **H**. A rescue time course was conducted with *osr1* MO and *osr1* cRNA to determine when the *osr1* cRNA dosage became insufficient to rescue. While there was a 90% rescue rate at 24 hpf, by 48 hpf, this rate had dropped to 20%. This indicated that continued *osr1* is needed for normal tubule development. P-values: **p<0.001, N.S. = not significant. For tubule rescue at 24 hpf, p-values were obtained from arcsin transformed kidney to body percentage calculations for each group. Scale bar is 50 µm for all images.

We also conducted a rescue time course by co-injecting *osr1* MO and *osr1* cRNA and performed WISH using *cdh17* to assess the tubule during a number of stages. While 90% of animals injected with both constructs exhibited a WT tubule length at 24-27 ss, by 36 hpf, only 50% showed a rescue (Figure 3H, Figure 3 - figure supplement 1B). At 48 hpf, only 20% of injected embryos had a WT length pronephric tubules while 80% had a unilateral or bilateral reduction (Figure 3H, Figure 3 - figure supplement 1B). Together, this indicated that the pronephros requires a continued presence of *osr1* in order for the tubule population to be maintained as development progressed.

### *wnt2ba* is a novel podocyte marker and regulator

Given the importance of *osr1* to podocyte development and maintenance, we wanted to identify downstream factors that promote podocytes. It was previously shown that the canonical Wnt ligand *wingless-type MMTV integration site family, member 2Ba* (*wnt2ba*) is expressed in a similar proximal swath of IM as *osr1* (Neto et al. 2012). We observed a similar expression pattern of *wnt2ba* in the anterior IM as early as 13 ss (Figure 4A). To specifically determine which cells *wnt2ba* was expressed in, we conducted FISH studies. At the 20-22 ss, *wnt2ba* transcripts were colocalized in cells within the anterior most region of *pax2a+* and *wt1b+* IM (Figure 4B, Figure 4 - figure supplement 1A). At 15 ss, *wnt2ba* transcripts were also colocalized with *wt1a/b+* podocyte progenitor cells (Figure 4 - figure supplement 1A,B). At 24 hpf, *wnt2ba* was expressed in both *wt1b*+ podocyte precursor cells and in neighboring cells of the IM (Figure 4C). By 48 hpf, *wnt2ba* was restricted to the podocytes and overlapped precisely with *wt1b* expression (Figure 4C). Taken together, we conclude that *wnt2ba* is a novel podocyte marker. We also examined the expression of the zebrafish *wnt2ba* paralogue, *wnt2bb*, at 24 hpf using FISH, but determined that transcripts were located anterior to the podocyte and kidney fields (Figure 4 - figure supplement 2).

**Figure 4:**
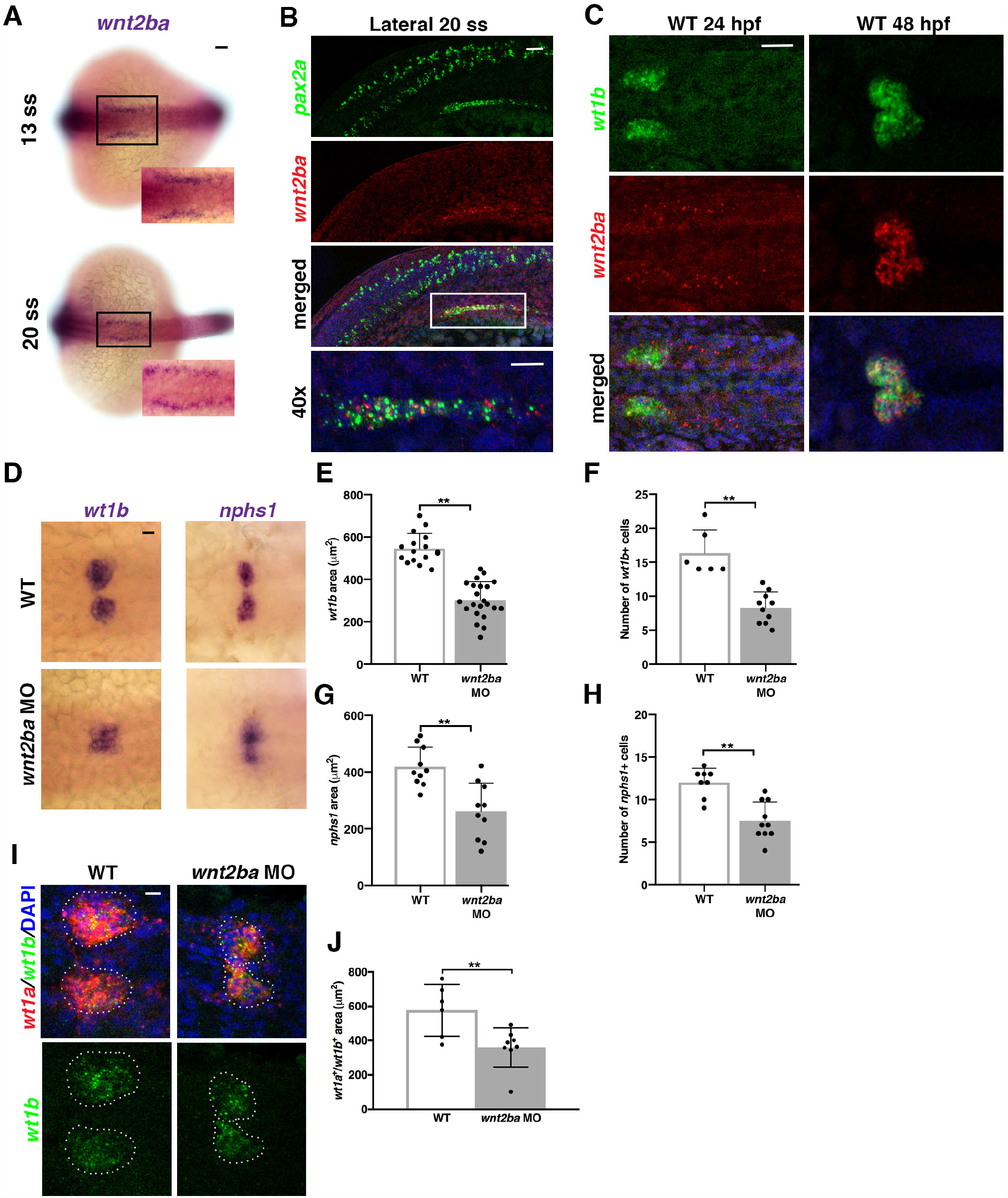
*wnt2ba* is a podocyte marker and regulator. **A**. *wnt2ba* is expressed in bilateral stripes as early as 13 ss. Scale bar is 30 µm. **B**. *wnt2ba* (red) is expressed within the anteriormost region of the IM, as shown by co-localization with *pax2a* (green). White box denotes area of co-localization, which is magnified in bottom panel. DAPI (blue) marks nuclei. Scale bar is 15 µm. **C**. *wnt2ba* (red) also colocalized with the podocyte marker *wt1b* at 24 hpf, though at this time point it was also expressed in putative neck segment domain. By 48 hpf, the *wnt2ba* domain was specified to the podocytes. Scale bar is 30 µm. **D-H**. Podocyte area and cell number was assessed in *wnt2ba* morpholino-injected animals and determined to be reduced compared to WT controls. Both *wt1b* and *nphs1* showed a significant decrease in domain area in *wnt2ba* morphants compared to WT embryos. **I**,**J**. FISH with *wt1a* and *wt1b* was performed at 24 hpf. There was a significant area reduction in podocyte domain seen in *wnt2ba* morphants compared to WTs. Scale bar is 10 µm. P-values: **p<0.001, *p<0.05, N.S. = not significant. Scale bar is 30 µm.

Given its expression in podocyte progenitors, we hypothesized that Wnt2ba might have roles in podocyte specification or differentiation. To explore whether *wnt2ba* is needed for proper podocyte formation, we performed *wnt2ba* loss of function studies. We first verified a morpholino that blocked splicing at exon 1, as well as a morpholino that targeted the start site (Figure 4 – figure supplement 3). When *wnt2ba* morphants were examined at 24 hpf, there was a significant reduction in the expression of *wt1b* and *nphs1* that corresponded to a smaller podocyte area and net cell number. (Figure 4D-H). We also found that the area of *wt1a+/wt1b+* coexpressing podocytes was decreased in *wnt2ba* morphants at 24 hpf (Figure 4I,J). Furthermore, the decrease in podocyte number in *wnt2ba* morphants occurred between the 15 and 22 ss, suggesting *wnt2ba* is required to maintain the podocyte lineage (Figure 4 – figure supplement 4). In contrast, *wnt2ba* morphants showed no discernable changes in development or maintenance of the *cdh17*+ nephron tubule (Figure 4 - figure supplement 3E). Collectively, these data lead us to conclude that *wnt2ba* is a significant regulator of podocyte ontogeny.

### *osr1* promotes *wnt2ba* in the podocyte developmental pathway

Previous research has demonstrated that *osr1* morphants exhibited a dramatic decrease in the *wnt2ba* pronephric domain, though *wnt2ba* morphants had no notable change in *osr1* expression (Neto et al. 2012). They postulated that *osr1* acts to promote *wnt2ba* in the IM which allows for proper pectoral fin development to occur (Neto et al. 2012). This led us to hypothesize that this same genetic cascade in the IM promotes the formation of proximal pronephric tissues, such as the podocytes, and is dysfunctional in *ocn-/-*.

In congruence with this prior study, we found that *wnt2ba* is significantly reduced in *ocn-/-*at both 15 ss, and is almost completely absent by 24 hpf (Figure 5A). We also observed that *wnt2ba* and *osr1* transcripts were colocalized in a population of presumptive IM cells at 15 ss and 22 ss, putting them in the right place and the right time to interact (Figure 5B, Figure 5 - figure supplement 1). Overexpression of *wnt2ba* led to an increase in podocyte number and domain area in injected WT embryos, as seen with an increase in the markers *wt1b* and *nphs1* (Figure 5C-H). While injection with *osr1* MO alone leads to diminished podocytes, co-injection of *wnt2ba* cRNA with *osr1* MO led to a rescue in podocyte area and cell count (Figure 5C-H). Together, this indicates that *wnt2ba* is sufficient to drive podocyte development, and does so downstream of *osr1*.

**Figure 5:**
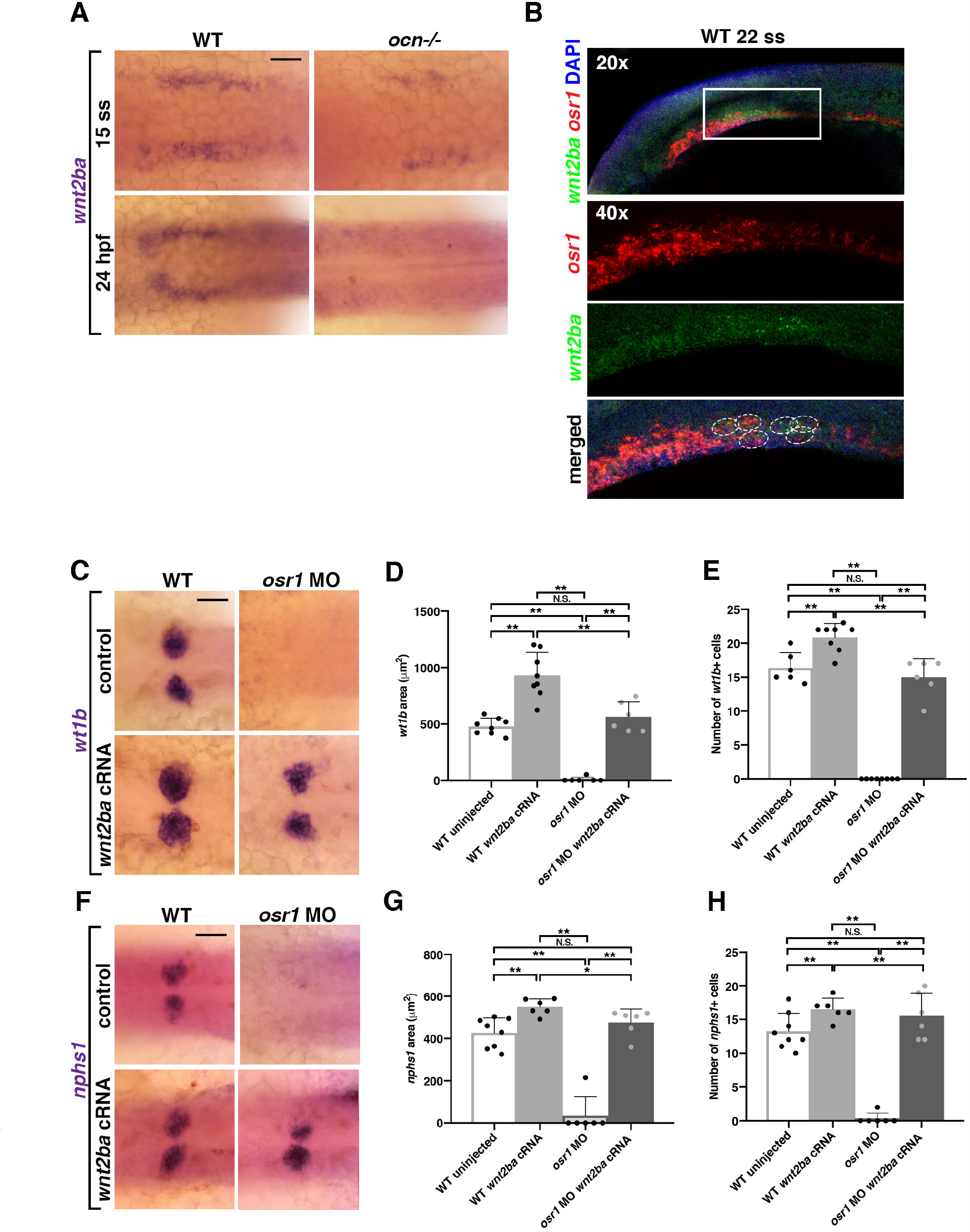
*wnt2ba* is sufficient for podocyte development downstream of *osr1*. **A**. When *wnt2ba* was assessed at 15 ss, in *ocn-/-* and WT siblings, it was evident that staining was reduced in mutants. By 24 hpf, *wnt2ba* staining was almost completely absent, tracking with the loss of other podocyte markers in *ocn-/-*. Scale bar is 30 µm. **B**. FISH experiments showed that *osr1* and *wnt2ba* colocalized in cells at 22 ss. **C-H**. Embryos were injected with *osr1* MO and/or *wnt2ba* cRNA at the one cell stage and podocytes and the developing slit diaphragm were visualized at 24 hpf using *wt1b* and *nphs1*, respectively. Embryos injected with *osr1* MO alone showed few podocyte or slit diaphragm cells, while embryos injected with *wnt2ba* cRNA alone had an increased podocyte area. Co-injected embryos had a partial rescue of podocytes, indicating that *wnt2ba* is a downstream factor in the podocyte pathway. A minimum of 5 individuals were imaged for quantification. P-values: **p<0.001, *p<0.05, N.S. = not significant. Scale bar is 30 µm.

### *hand2* suppresses podocyte development by restricting *wnt2ba* expression and podocyte development

The bHLH transcription factor *heart and neural crest derivatives expressed 2* (*hand2*) has been shown to be antagonistic to *osr1* in early mesoderm development (Perens et al. 2016). The loss of *osr1* leads to decreases in podocytes and tubules and an increase in hemangioblasts; in contrast, loss of *hand2* results in expansions in renal cells at the expense of vasculature (Perens et al. 2016). Concomitant knockdown of *osr1* and *hand2* rescues tubule development (Perens et al. 2016) and podocyte development (Perens et al. 2020, bioRxiv).

When we knocked down *hand2* using an ATG morpholino, we observed a separation in the *myosin light chain 7* (*myl7*) heart field at 22 ss that matched previously observed phenotypes (Maves et al. (2009) (Figure 6 - figure supplement 1). Knockdown of *hand2* also caused a significant increase in *wt1b*+ podocyte domain area and cell number (Figure 6A-C). While uninjected *ocn-/-*embryos had little to no podocytes, injecting *ocn-/-*with *hand2* MO resulted in an expansion in podocyte number and area that was significantly different from mutants (Figure 6A-C). Similarly, *hand2* morphants had a significantly larger *wnt2ba* domain, and *hand2/osr1* MO coinjection rescued the usually abrogated *wnt2ba* domain (Figure 6D,E). This indicated that imbalance of *hand2* and *osr1* leads to changes in *wnt2ba* expression, which alters podocyte development.

**Figure 6:**
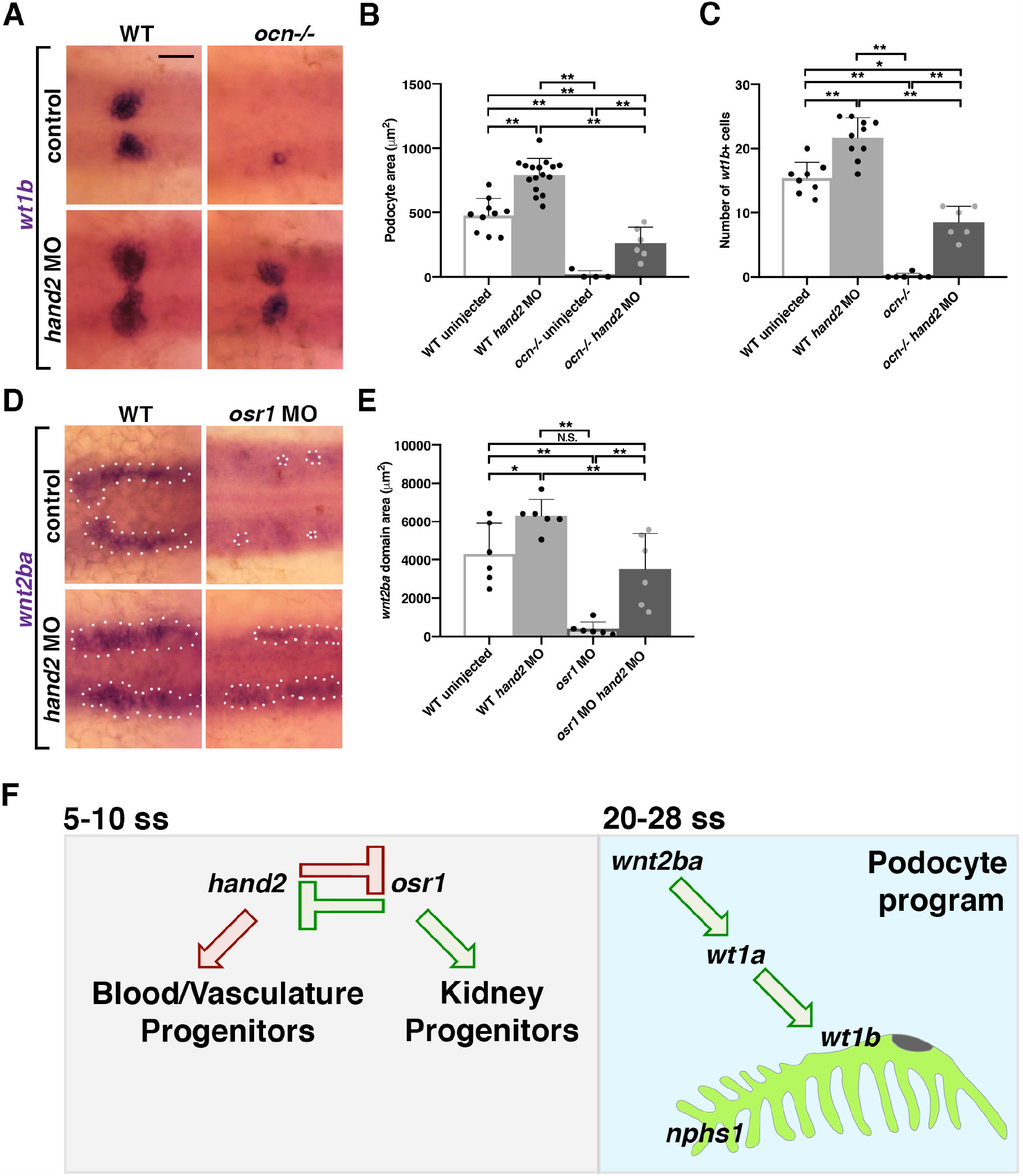
Acting in opposition to *osr1, hand2* inhibits *wnt2ba*-driven podocyte development. **A**,**B**,**C**. Embryos from *ocn* incrosses were injected with *hand2* MO. Podocytes area and cell number were partially rescued in *ocn-/-*injected embryos, and were expanded in WT injected siblings. This signified that *hand2* suppresses podocyte formation. **D**,**E**. Similarly, *wnt2ba* expression was rescued in *osr1/hand2* morphants compared to uninjected WT controls. Injection of *hand2* MO alone led to a significant increase in the *wnt2ba* domain. All images were at 24 hpf, scale bar is 30 µm. A minimum of 5 individuals were imaged for quantification. P-values: **p<0.001, *p<0.05, NS = not significant. **F**. *osr1* promotes podocyte and kidney lineages and suppress blood and vasculature, while *hand2* acts in opposition. Imbalance of either of these factors leads to changes in mesodermal fates. Podocyte development is one example of a mesodermal fate that is altered by imbalance of *osr1/hand2*. This is because the downstream factor *wnt2ba* is decreased without *osr1*, yet increased in the absence of *hand2. wnt2ba* endorses the podocyte factors *wt1a/b*, which have been shown to be required for formation of podocytes and the slit diaphragm (*nphs1*).

## DISCUSSION

While there are dozens to hundreds of podocyte diseases and maladies that have been characterized, the genetic explanations for their origin and progression is lacking. One reason for this is that there are relatively few factors that are known to promote the development of these specialized epithelia. Continuing to identify these factors is critical for future diagnostics and treatments for podocytopathy. In this study, we have both re-examined a previous factor shown to promote podocyte fates, *osr1*, and identified a new downstream regulator, *wnt2ba*.

*ocn* was identified as a mutant of interest in a forward genetic screen due to displaying pericardial edema and decreases in podocytes and proximal tubule. Through a whole genome sequencing approach, we determined that *ocn-/-*harbors a SNP in exon 2 that leads to a premature stop in *osr1*. This SNP was confirmed as the causative lesion in *ocn-/-*when *osr1* cRNA could rescue each of these phenotypes. Upon confirmation that *ocn* was an *osr1* mutant line, we next sought to fully assess how *osr1* loss of function impacts kidney development in the context of a zebrafish mutant. We found that *osr1* is needed to maintain renal progenitors and inhibit the development of hemangioblasts.

Further, we established a genetic pathway controlled by *osr1* that regulate podocyte survival by promoting *wnt2ba* expression. We found that *wnt2ba* is an IM/podocyte marker that is likewise diminished in *ocn-/-*. Loss and gain of *wnt2ba* leads to a decrease and increase in podocyte area, demonstrating that *wnt2ba* is both necessary and sufficient to drive podocyte development. Notably, *wnt2ba* can rescue podocytes in an *osr1-*deficient background, which places this factor downstream of *osr1* (Figure 6F). Finally, the *osr1/wnt2ba* podocyte pathway is negatively regulated by *hand2* (Figure 6F).

### *osr1* acts to promote podocytes

The earliest known podocyte marker in zebrafish is *wt1a*, though the paralogue, *wt1b*, that appears at 12 ss is expressed in a more specific territory (Bollig et al. 2006; Bollig et al. 2009). It has also been suggested that *wt1a* is more dominant than *wt1b*, as knockdown of *wt1a* leads to loss of *nphs1/2* while knockdown of *wt1b* causes less dramatic podocyte phenotypes (Perner et al. 2007). Zebrafish literature has shown that *osr1* morphants exhibit reductions in *wt1b, lhx1a*, and *nphs1/2* at 24 hpf that have also been observed in *ocn-/-*(Mudumana et al. 2008; Tomar et al. 2014, Tena et al. 2007). However, the relationship between *wt1a* and *osr1* has yet to be fully understood. Tomar et al. (2014) placed *wt1a* upstream of *osr1* due to *osr1* being reduced in *wt1* morpholino-injected embryos and *wt1a* expression being interpreted as “unchanged” in *osr1* morpholino-injected animals. However, in our studies, *osr1* morphants do exhibit alterations in *wt1a*+ cell organization and a restriction in domain that phenocopies *ocn-/-*. Mouse studies have shown that *WT1+/-;OSR1+/-*mice exhibit reduced kidneys, suggesting that these factors act cooperatively in kidney and podocyte development (Xu et al. 2016). If *osr1* and *wt1a* did have a similar synergistic relationship in zebrafish kidney development, this would also explain reports that *wt1* morphants exhibit a loss of podocytes and proximal tubules reminiscent of *osr1* loss of function models (Tomar et al. 2014). While there are currently limitations in using anti-Osr1 antibodies in any *in vivo* model, progress in this area is needed in order to ascertain if *wt1a* and *osr1* are directly interacting during kidney development.

### *osr1* is needed for kidney cell maintenance

While *ocn-/-*exhibit normal patterning of IM early in development, by the time specification to pronephros is beginning to occur around 15 ss, the anterior domain is absent. Our experiments demonstrated that this is due to two events; (1) an expansion of hemangioblasts, and (2) apoptosis of podocyte progenitors in this region. Work in chick and mouse has shown that while mesonephric tissues and markers are present, apoptosis occurs within nephrogenic mesenchyme that keeps metanephric tissues from forming in *Osr1* knockout animals (Wang et al. 2005; James et al. 2006). Further, previous studies have shown that *Osr1* acts synergistically with factors such as *Wt1* and *Six2* to renew renal stem cell pools to inhibit premature differentiation and thus cell death (Xu et al. 2014). A similar apoptosis event has not been recorded in *osr1* loss of function zebrafish models prior to this study, and we hypothesize that *osr1* plays a similar role in progenitor self-renewal in zebrafish.

The expansion in hemangioblast domain in *osr1* morphants has been documented by other groups, where it was suggested that *pax2a*+ cells were converting to *tal*+ cells (Mudumana et al. 2008). Additionally, the expansion in vessel progenitors has been reported in an *osr1* TALEN mutant (Perens et al. 2020, bioRxiv). However, our results add one further element to these early events, as we have captured cell apoptosis in *pax2a+* cells of *osr1* mutant embryos. Further, our studies have revealed that the timing of the *pax2a* domain decrease and hemangioblast domain increase is not equivalent. The hemagioblasts expand hours prior to the loss of the anterior IM domain. We postulate that *osr1* may inhibit hemangioblast formation either indirectly or in an independent mechanism than it uses to promote IM and podocytes.

### *wnt2ba* is a novel regulator of podocyte development

*wnt2ba* is a ligand that functions in the canonical Wnt/beta-catenin pathway. As a member of this pathway, *wnt2ba* acts to promote cell growth, differentiation and migration during development. In regards to kidney development, it has been shown that *Wnt2b* can be detected in the kidney stroma in mice as early as E11.5 (Lin et al. 2001; Iglesias et al. 2007), and in humans WNT2B is expressed in fetal kidney stroma (Combes et al., 2019). In addition, cells expressing Wnt2b promote ureteric branching in culture (Lin et al. 2001). *Wnt2/2b* is also paramount to normal lung and pectoral fin development in both aquatic and mammalian species (Goss et al. 2009, Neto et al. 2012). Interestingly, *osr1* has been shown to act downstream of retinoic acid signaling yet upstream of *wnt2b* in both pectoral fin development in zebrafish (Neto et al. 2012) and in lung progenitor specification in foregut endoderm in *Xenopus* (Rankin et al. 2012). However, our study has both evaluated the role of *wnt2ba* as a regulator of kidney development and placed its function downstream of *osr1* to specifically promote the podocyte lineage. Further, we show that *osr1* promotes *wnt2ba* expression during podocyte development through a mechanism involving the inhibition of *hand2*. In synchrony with our data, a recent report similarly found that reciprocal antagonism between *osr1* and *hand2* is essential for the normal emergence of *wt1b+* podocyte precursors (Perens et al. 2020, bioRxiv).

We show in the present study that *wnt2ba* is a regulator of podocyte development, but that loss of *wnt2ba* does not cause compelling changes in PCT or tubule length. Another study by Lyons et al. (2009) showed that broad inhibition of Wnt signaling through heat-shock activation of *dkk1* lead to an abrogation in the zebrafish pronephros that resembles *osr1* loss of function models. Wnt ligands are highly regionalized to allow for precise regulation during tissue development (Iglesias 2007; Verkade and Heath 2008). Our findings that *wnt2ba* is restricted to the podocytes by 48 hpf could reflect regional specificity. This suggests that there are other Wnt ligands and receptors that act to regulate certain kidney lineages in zebrafish development. Loss of one or more of these factors in combination with *wnt2ba* could lead to an anterior truncation of the pronephros that resembles the experiments from Lyons et al. (2009). Future studies are needed to discern these factors and additional downstream targets of both *wnt2ba* and *osr1*.

Taken together, these results have allowed us to garner new insights into podocyte development in zebrafish. By selecting *ocn* as a mutant of interest from our ENU screen, we have discovered an *osr1* mutant and confirmed its significance in zebrafish pronephros development in an unbiased manner. We have expanded on these previous findings by demonstrating that *osr1* is required for to inhibit apoptosis in specified kidney precursors, and later for nephron cell maintenance. We have also ascertained new roles for *osr1* in promoting *wnt2ba* expression, which it does in part through antagonism of *hand2*. Finally, our results show that *wnt2ba* mitigates podocyte development downstream of the *osr1/hand2* interaction. Given how little is known about CAKUT and kidney agenesis, findings from genetics studies such as the present work are crucial to furthering our understanding about the causes and solutions to these disease states.

## METHODS

### Creation and maintenance of zebrafish lines

Zebrafish were housed in the Center for Zebrafish Research in the Freimann Life Science Center at the University of Notre Dame. All experiments and protocols used in this study were approved by the Institutional Animal Care and Use Committee (IACUC). We performed an ENU haploid genetic screen as described in Kroeger et al. (2017) and Chambers et al. (2018).

### Live imaging and dextran injections

Embryos were grown in E3 media at approximately 28°C. For live imaging, embryos were placed in a solution of 2% methylcellulose/E3 and 0.02% tricaine and placed in a glass depression slide. For dextran injection experiments, embryos were also incubated with 0.0003% phenylthiourea (PTU) in E3 to inhibit pigment development. At 3 dpf, *ocn*::*cdh17*::GFP animals were anesthetized and injected with 40 kDA rhodamine-dextran. Embryos were then examined and imaged 24 hours after injection.

### WISH, FISH, IF, sectioning and image acquisition

WISH was performed as described in previous studies (Wingert et al. 2007, Wingert and Davidson 2011, Li et al. 2014, Cheng and Wingert 2015). For each marker, embryos from at least 3 sets of parents were assessed and 5 mutants and 5 siblings were imaged. Immunocytochemistry (ICC) was performed as previously described (Kroeger et al. 2017, Marra et al. 2018). Embryos from WISH experiments were embedded in JB-4 plastic blocks and cut to obtain 4 µm sections which were counterstained with methylene blue (0.5%) to mark nuclei. Alcian blue staining was performed as described (Chambers et al. 2018).

### Genotyping

Direct genotyping on *ocn* fin clips and embryos was carried out by PCR amplification of exon 2 of *osr1* (Forward primer: CCCCATTCACTTTGCCACGCTGCACCTTTTC, reverse primer: CTGTGGTCTCTCAGGTGGTCCTGCCTCCTAAA). Dilutions of purified PCR products were then subjected to Sanger sequencing by the Genomics Core at University of Notre Dame using the forward primer.

### Morpholinos and RT-PCR

*osr1* morpholino (ATCTCATCCTTACCTGTGGTCTCTC) was first described in Mudumana et al. (2008) and was designed to block the splice donor site of exon 2. We used the primers GTGACTGTATCTGAATCCTCTTATTTTGGATCGTCTCGCTTCACAAAGAACTG, and CTGTAGGCTATGGAAGTTTGCCTTTTCAGGAAGCTCTTTGGTCAG to perform RT-PCR as described in (Kroeger et al. 2017) to confirm the interruption of exon 2 splicing activity. *wnt2ba* splice blocking morpholino (CTGCAGAAACAAACAGACAATTAAG) was previously utilized in Neto et al. (2012). However, we did not find their methods of RT-PCR to be specific to *wnt2ba* transcripts, so we designed RT-PCR primers to amplify the entire transcript; (Forward primer: ATGCCAGAGTGTGATGGAGTTGGGTGCGCGTCGCCGGCGC, (Reverse primer: GCTGGAGCGAGACCACACTGTGTTCGGCCGC), and additionally looked for the presence of intronic sequence with (Intronic forward primer: ATCACAGGGGTATCATTATCACAAAAAATTGTAAATAAATG). While this splice blocking morpholino was the primary method of *wnt2ba* knockdown, a *wnt2ba* ATG morpholino (ACCCAACTCCATCACACTCTGGCAT, Wakahara et al. 2007) was used to confirm phenotypes seen with the splicing morpholino. The *hand2* ATG morpholino (CCTCCAACTAAACTCATGGCGACAG) was used as described in Maves et al. (2009).

### Statistics and measurements

Absolute domain lengths and area measurements were taken from five representative embryos per treatment using Fiji is just ImageJ. Averages, standard deviation, and unpaired student’s t-tests were performed in Microsoft Xcel and GraphPad Prism. In experiments where *osr1* cRNA was used, body axis measurements were taken for injected and uninjected embryos. The tubule measurements for each group was divided by the body length measurement to discern what percentage of the body length was occupied by the kidney. To normalize the data, these percentages were subjected to arcsine degree transformation and then run through a student’s t-test to determine significance.

## Abbreviations

(CKD): chronic kidney disease
(ENU): N-ethyl-N-nitrosourea
(dpf): days post fertilization
(hpf): hours post fertilization
(WISH): whole mount *in situ* hybridization
(FISH): flourescent *in situ* hybridization
(IF): immunofluorescence
(ICC): immunocytochemistry
(IM): intermediate mesoderm
(CAKUT): congenital anomalies of the kidney and urinary tract
(aa): amino acid
(ZIRC): Zebrafish International Research Center
(cRNA): capped RNA
(ss): somite stage
(WT): wild-type
(*lhx1a*): *LIM homeobox 1a*
(*pax2a*): *paired box 2a*
(*wt1a*): *wilms tumor 1a*
(*wt1b*): *wilms tumor 1b*
*(nephrin) (nphs1)*: *nephrosis 1, congenital Finnish type*
*(podocin)*(*nphs2*): *nephrosis 2, idiopathic, steroid-resistant*
(*osr1*): *odd-skipped related transcription factor 1*
(*cdh17*): *cadherin 17*
(*osr2*): *odd-skipped related transcription factor 2*
(*wnt2ba*): *wingless-type MMTV integration site family, member 2Ba*
(*wnt2bb*): *wingless-type MMTV integration site family, member 2Bb*
(*hand2*): *heart and neural crest derivatives expressed 2*
(*myl7*): *myosin light chain*

## Supplemental Figure Legends

**Figure 1, Supplement 1:**
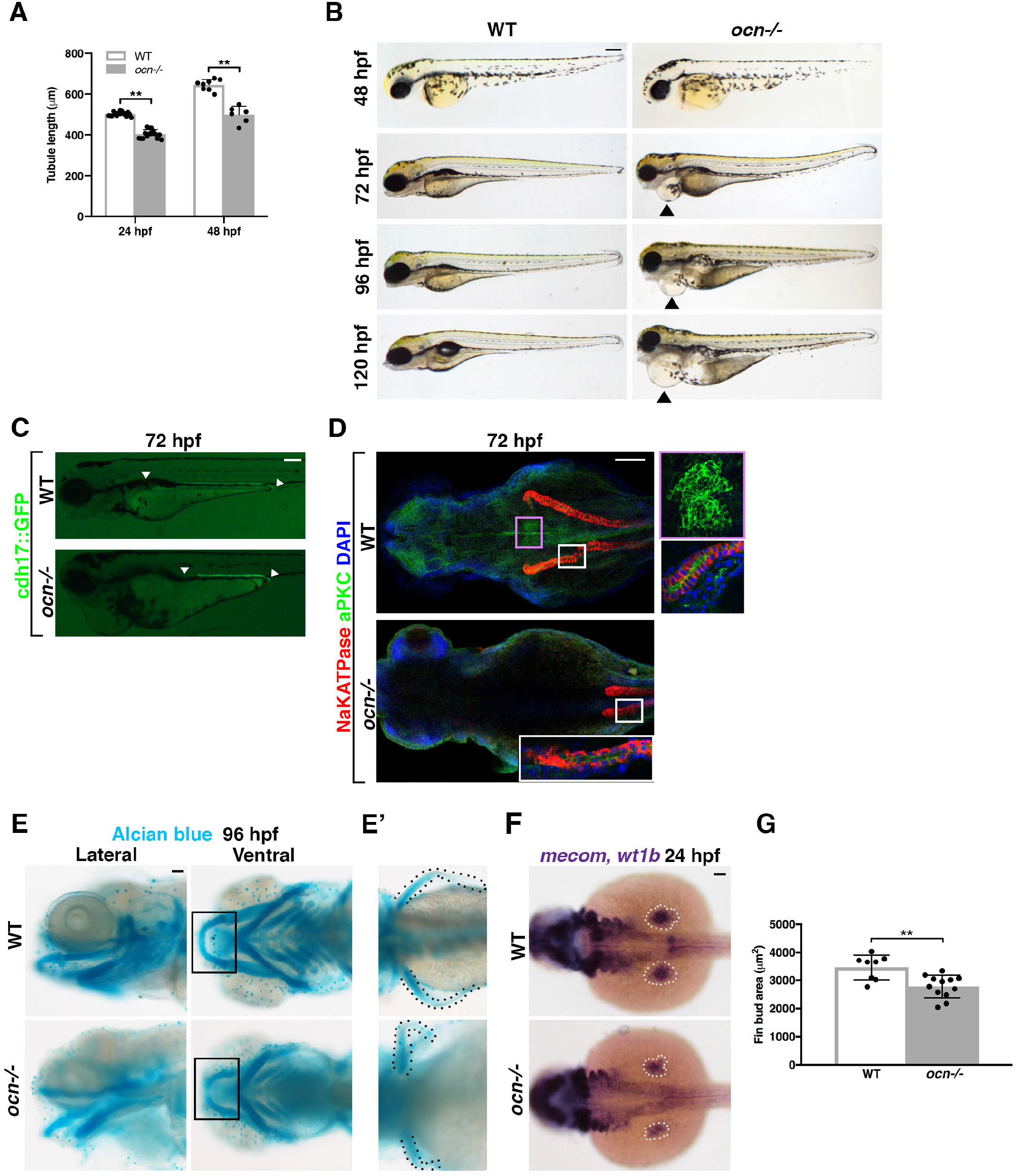
Additional *ocn* phenotypes. **A**. Conducting absolute length measurements of *cdh17* tubule revealed that *ocn* have significantly shorter nephrons tubules at 24 hpf and 48 hpf compared to WT siblings. **B**. A live time course of *ocn-/-*revealed pericardial edema beginning at 72 hpf, as indicated by black arrow heads. Scale bar is 70 µm. **C**. *ocn+/-*were crossed with a cdh17::GFP transgenic line to create *ocn* carriers with GFP-labeled pronephric tubules. Pictured here are 72 hpf cdh17::GFP embryos and F2 *ocn-/-*::cdh17::GFP embryos. White arrowheads demarcate the start and terminus of the pronephros, and it is evident that mutants have a shorter pronephros. Scale bar is 30 µm. **D**. Similarly to WISH results with *cdh17* and *wt1b*, IF revealed that *ocn-/-*embryos had a truncation in NaKATPase+ tubule and absence of aPKC+ glomerulus. **E**. The acidic stain, alcian blue, was used to show cartilage in developing *ocn-/-* and WT siblings at 96 hpf. Lateral and ventral views showed that the jaw developed in an improper orientation with unfused Meckel’s cartilage (box). Scale bar is 30 µm. Pectoral fins in *ocn-/-*mutants were also malformed. **F**,**G**. The pectoral fins arise from the fin buds (white dotted area), which are marked by *mecom* at 24 hpf. Fin bud area measurements were decreased in mutants, which could be genotyped by the absence of *wt1b*+ podocytes. Scale bar is 30 µm. P-values: **p<0.001.

**Figure 1, Supplement 2:**
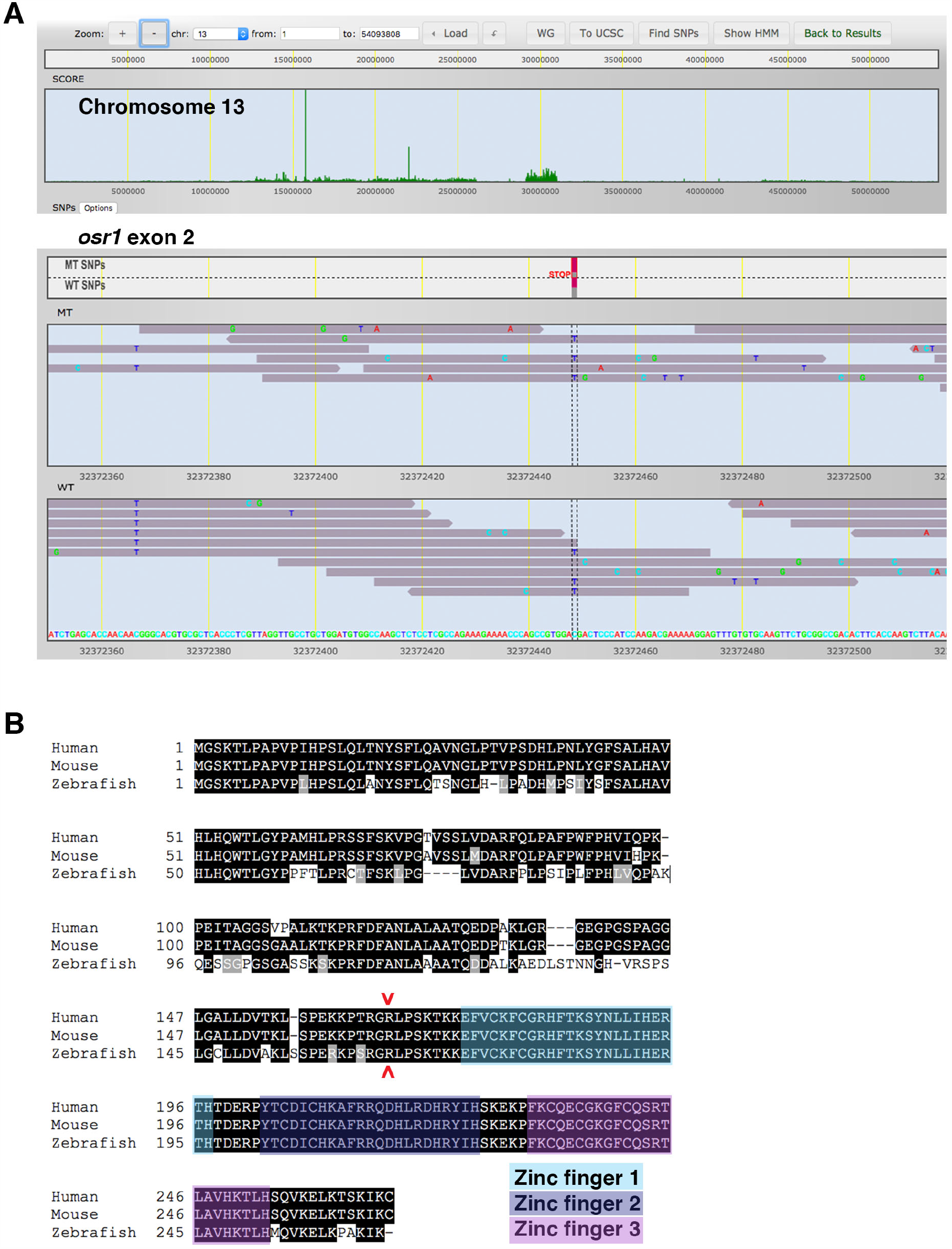
A genetic lesion in the *osr1* transcription factor would result in truncated osr1 protein in the *ocn* mutant line A. Whole genome sequencing was conducted on 24 hpf *ocn* mutants and WT siblings. SNPtrack analysis indicated that the genetic lesion responsible for the *ocn-/-*phenotypes was most likely located on chromosome 13. Specifically, a C to T missense mutation in exon 2 of *osr1* would cause a premature stop codon. **B**. Zebrafish, mouse and human OSR1 protein contains three zinc-finger binding domains. The predicted SNP would result in a substitution from an arginine to a stop codon before the transcription of the zinc-finger binding domains, which are each 100% conserved across zebrafish, mice and human OSR1 protein.

**Figure 1, Supplement 3:**
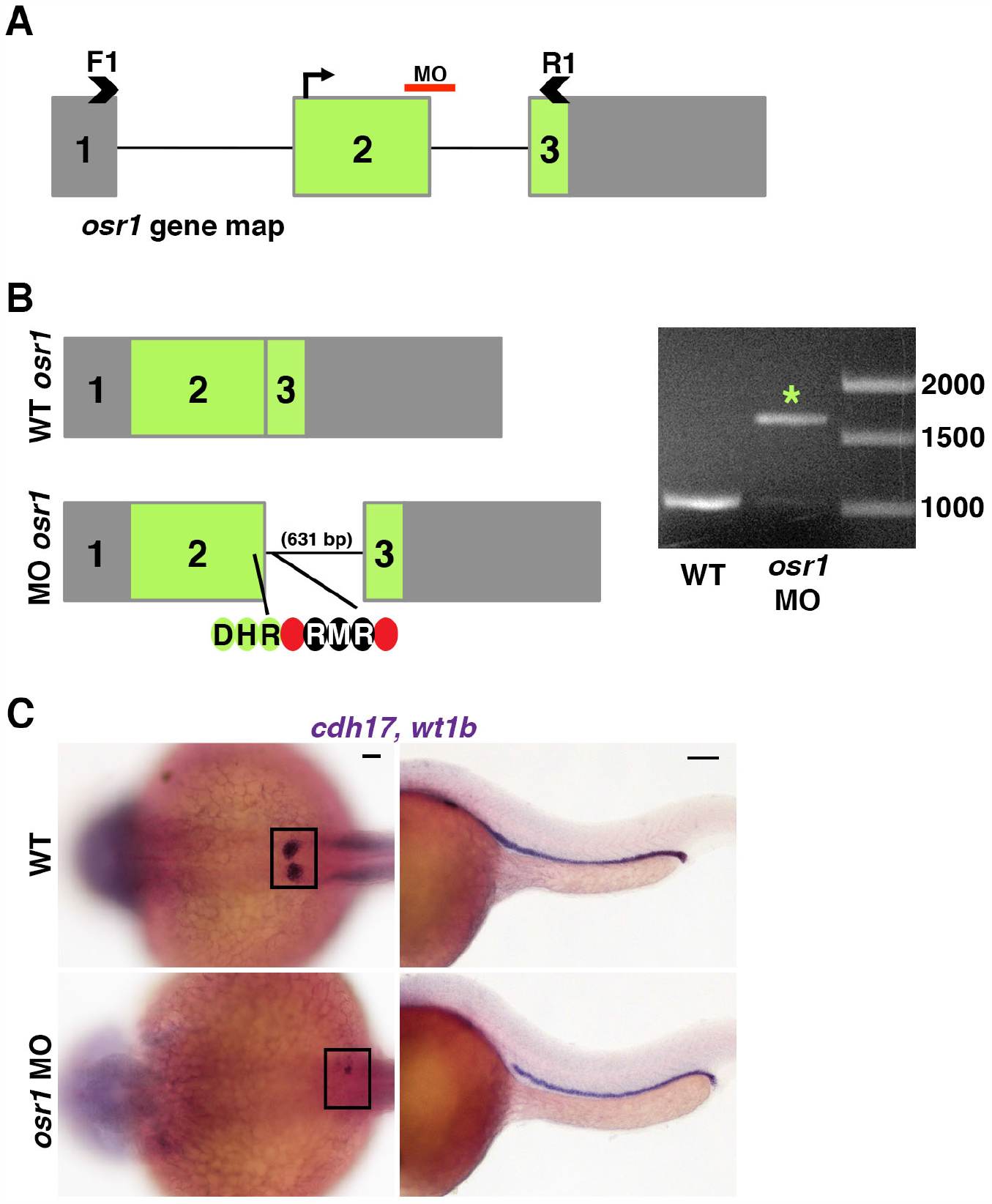
Verification of *osr1* splice-blocking morpholino phenotypes. **A**,**B**. A previously published morpholino was obtained to block splicing activity in exon 2 of *osr1*. Primers flanking exon 2 (arrowheads) were used to conduct RT-PCR to assess splicing activity in morphants at 24 hpf. In WTs with correct splicing, a 1000 bp product is obtained. In *osr1* morphants, intron 2 fails to be spliced out the gene, leading to a 631 bp increase in product. Further, the intronic sequence retained in *osr1* morphants contains in-frame stop codons. **C**. *osr1* morphants had dramatically decreased podocytes and a cropped pronephric tubule which resembled both previously published results and *ocn-/-*. Scale bar is 30 µm.

**Figure 2, Supplement 1:**
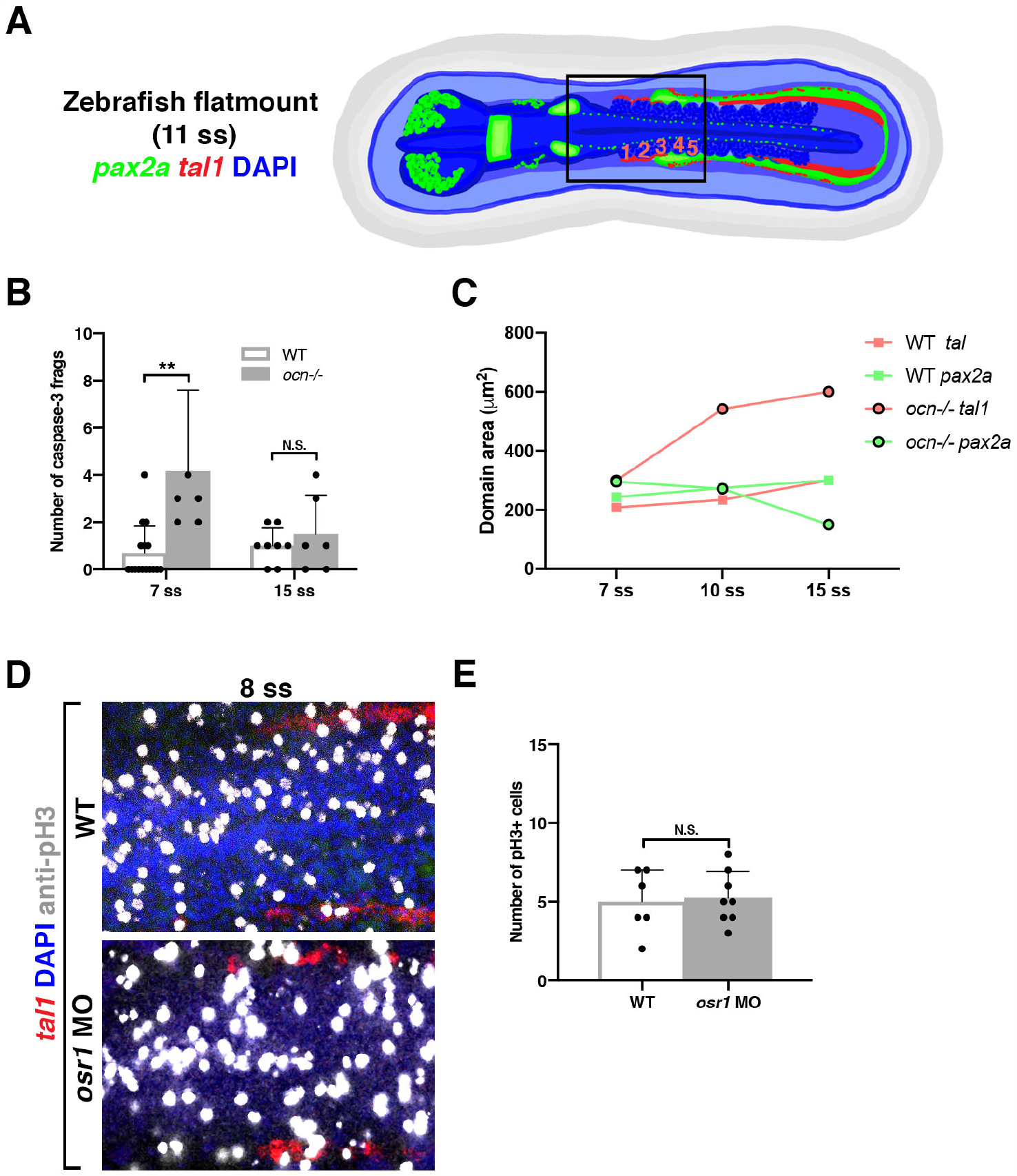
Additional analysis of cell dynamics in *osr1* deficient models. **A**. FISH experiments were conducted on young embryos (11 ss = 15 hpf) and flatmounted to be imaged. Probes for *pax2a* (green) were used to mark IM, and *tal1* (red), to mark hemangioblasts. DAPI (blue) marks nuclei and was used to distinguish cellular features such as muscular units known as somites. Counting somites allowed for accurate staging of embryos and a consistent location for tissue assessment. Areas and cell counts of mesodermal tissues were assessed from somites 1-5, as shown in the boxed area. **B**. There was a significant increase in caspase-3+ fragments in the *tal1/pax2a* area of interest in *ocn-/-*compared to siblings at 7 ss. However, by 15 ss, the number of caspase-3+ fragments in *ocn-/-*had returned to a WT level. **C**. Compared to WT sibings, the *tal1* domain steadily increased in *ocn-/-*from 7 to 15 ss. In contrast, the *pax2a* domain only became significantly smaller in *ocn-/-*mutants at the 15 ss. **D**,**E**. There was no significant change in the number of pH3+ cells in *osr1* morphants in the *tal1* domain between somites 1-5 at the 8 ss.

**Figure 3, Supplement 1:**
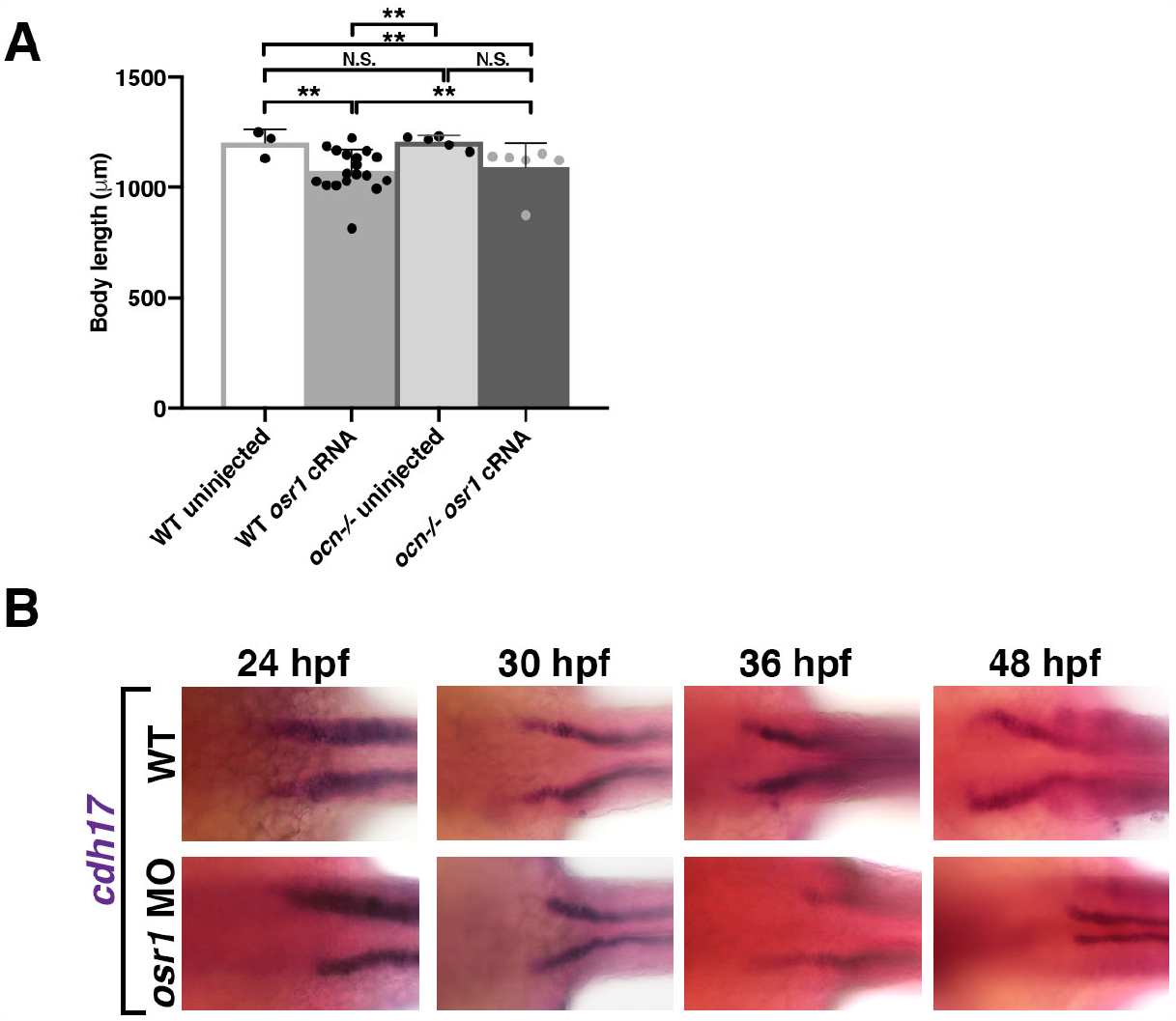
*osr1* cRNA injection additional analysis. **A**. *ocn-/-* and WT sibling embryos injected with *osr1* cRNA had significantly shorter body lengths than uninjected *ocn-/-*mutants and siblings. P-values: **p<0.001, N.S. = not significant. P-values were obtained by arcsin transforming percentages to normalize data. **B**. Time course of *cdh17* expression in WT and *osr1* MO injected animals. Scale bar is 20 µm.

**Figure 4, Supplement 1:**
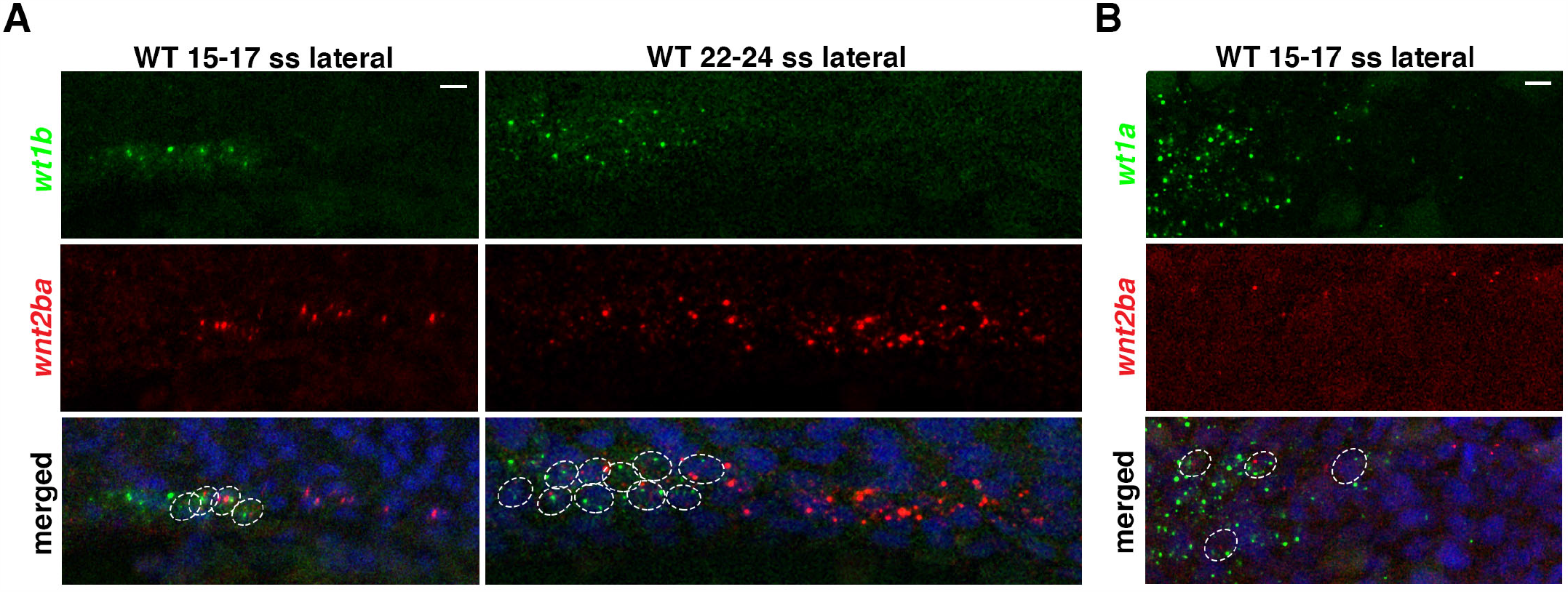
*wnt2ba* colocalizes with early, developing podocytes A,B. Lateral views of FISH experiments reveal that *wnt2ba* is present in developing *wt1a* and *wt1b* podocytes at 15-17 ss and 22-24 ss. Scale bar is 10 µm.

**Figure 4, Supplement 2:**
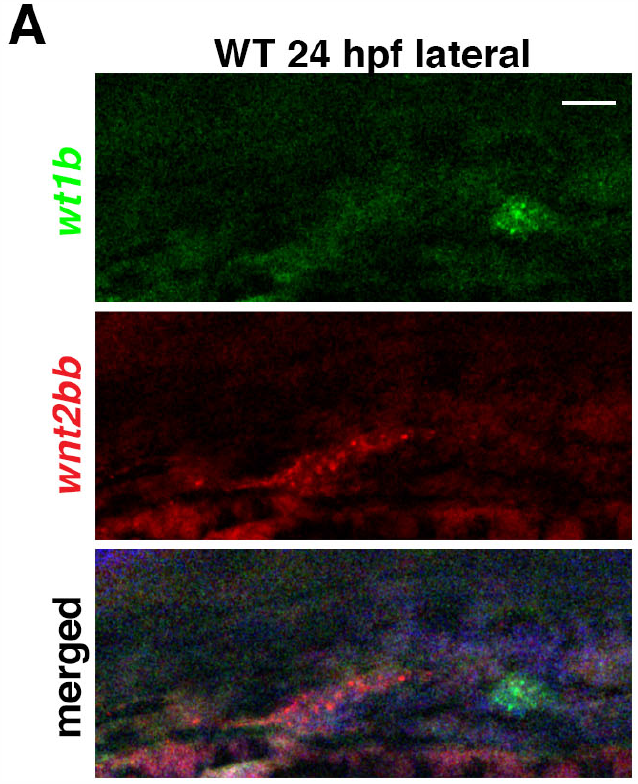
*wnt2bb* does not colocalize with podocytes. **A**. Lateral views of WT embryos stained using FISH reveal that *wnt2bb* is expressed anterior to *wt1b* podocytes at 24 hpf. Scale bar is 50 µm.

**Figure 4, Supplement 3:**
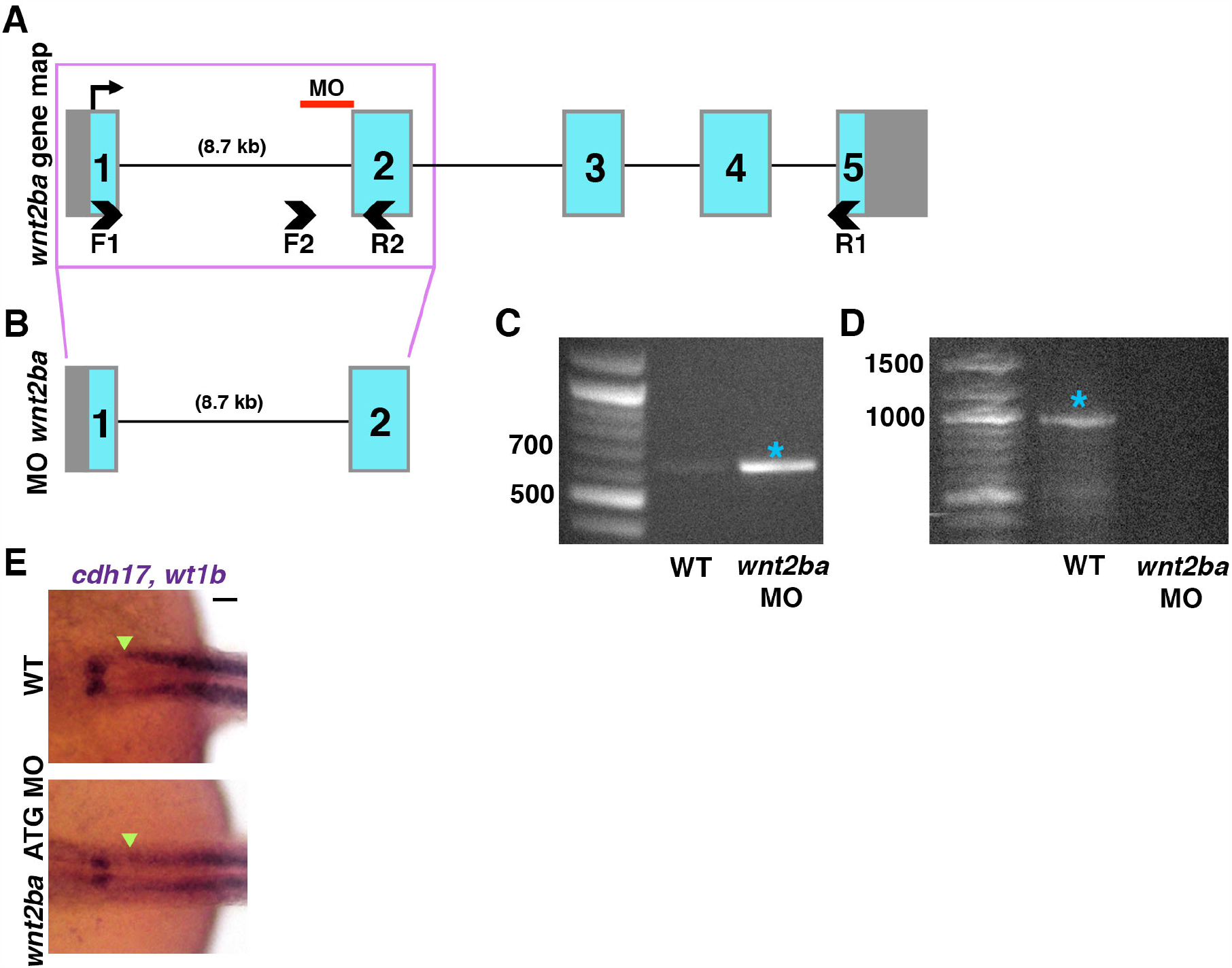
Assesment of *wnt2ba* MO through RT-PCR analysis. **A**. The gene *wnt2ba* contains 5 exons. A morpholino was utilized to block splicing activity in intron 1. **B**. In WT embryos, correct splicing activity occurs to splice exon 1 and 2 together, resulting in a ∼300 bp product as seen with F1/R1 primers. This product is greatly diminished in *wnt2ba* morphants, suggesting that correct splicing activity is limited. **C**. Primers were utilized that flanked the end of intron 1 and the middle of exon 2 in *wnt2ba* (F2/R2). In *wnt2ba* morphants, a strong band was present that indicated the presence of un-spliced, intronic sequence **D**. Primers flanking the entire ORF (F1/R2) were used as an additional metric to gauge splicing action. WTs exhibit a 1000 bp band which represents the *wnt2ba* ORF. *wnt2ba* morphants do not have this band, which further suggests that splicing activity is impaired. **E**. An ATG MO was used to knockdown *wnt2ba*. While podocytes were reduced using this reagent, the pronephric tubule was not affected. The start of the tubule is shown using green arrowheads. Scale bar is 50 µm.

**Figure 4, Supplement 4:**
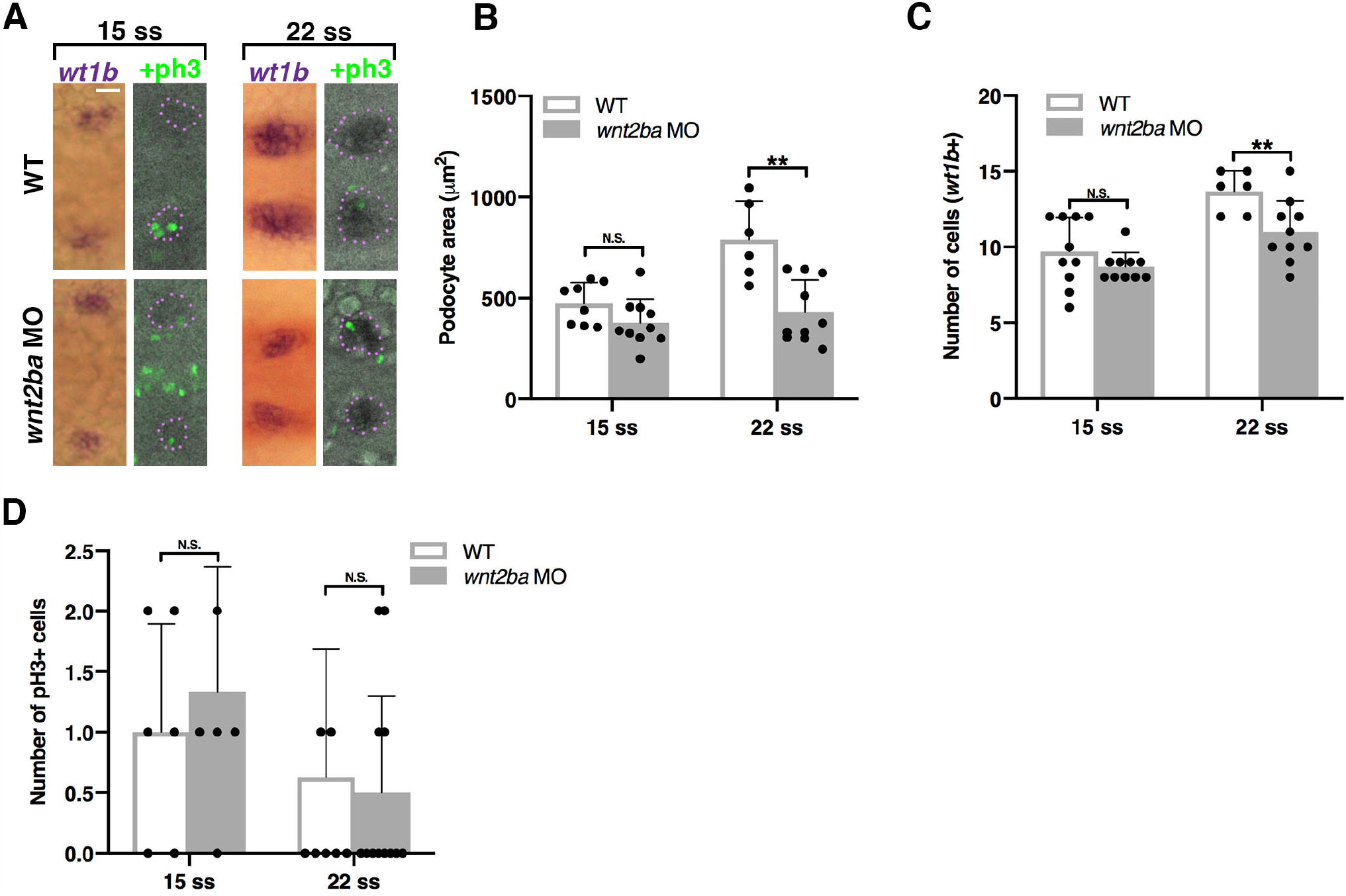
*wnt2ba* knockdown causes decreased podocytes at 22 ss. **A-D**. *wnt2ba* was knocked down using a splice-blocking MO and podocytes were examined by conducting WISH on 15 ss and 22 ss embryos using the marker *wt1b*. Anti-pH3 was used on these samples to assess proliferating cells. Podocyte area and cell counts were not different between morphants and WTs at 15 ss, though there was a significant decrease in podocyte cell number and domain area at 22 ss in morphants. However, there were no changes in the number of pH3+ cells in the *wt1b* area at either time point. Scale bar is 40 µm.

**Figure 5, Supplement 1:**
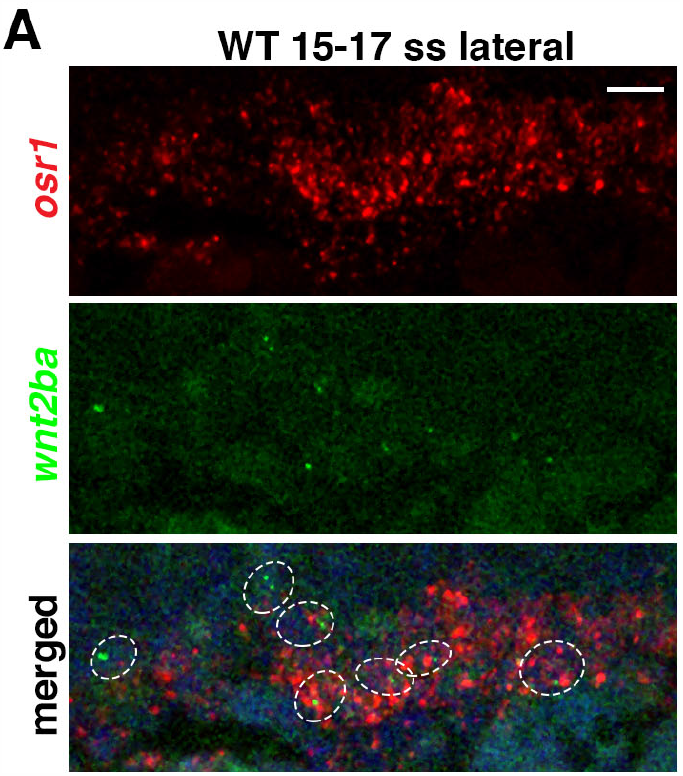
Additional stages of *wnt2ba* and *osr1* co-localization. **A**. In addition to 22 ss, it was also observed that *osr1* and *wnt2ba* colocalized in a subset of presumptive IM at 15-17 ss. Scale bar is 10 µm.

**Figure 6, Supplement 1:**
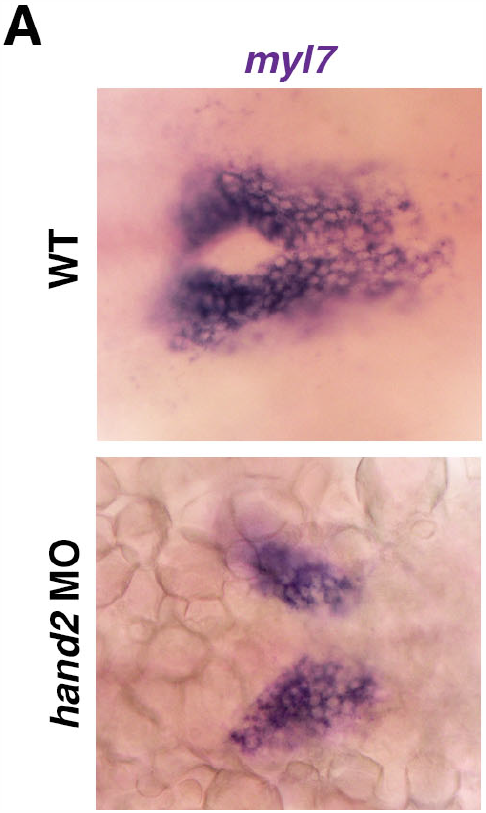
*hand2* MO replicates previous studies. **A**. The developing heart tube can be visualized with *myl7* at 22 ss. Embryos injected with *hand2* MO display cardia bifida, or a separation of heart precursors.

